# CPA-Perturb-seq: Multiplexed single-cell characterization of alternative polyadenylation regulators

**DOI:** 10.1101/2023.02.09.527751

**Authors:** Madeline H. Kowalski, Hans-Hermann Wessels, Johannes Linder, Saket Choudhary, Austin Hartman, Yuhan Hao, Isabella Mascio, Carol Dalgarno, Anshul Kundaje, Rahul Satija

**Affiliations:** New York Genome Center, New York, NY, USA; Center for Genomics and Systems Biology, New York University, New York, NY, USA; New York University Grossman School of Medicine, New York, NY, USA; Department of Genetics, Stanford University, Stanford USA; Department of Computer Science, Stanford University, Stanford USA

## Abstract

Most mammalian genes have multiple polyA sites, representing a substantial source of transcript diversity that is governed by the cleavage and polyadenylation (CPA) regulatory machinery. To better understand how these proteins govern polyA site choice we introduce CPA-Perturb-seq, a multiplexed perturbation screen dataset of 42 known CPA regulators with a 3’ scRNA-seq readout that enables transcriptome-wide inference of polyA site usage. We develop a statistical framework to specifically identify perturbation-dependent changes in intronic and tandem polyadenylation, and discover modules of co-regulated polyA sites exhibiting distinct functional properties. By training a multi-task deep neural network (APARENT-Perturb) on our dataset, we delineate a *cis*-regulatory code that predicts responsiveness to perturbation and reveals interactions between distinct regulatory complexes. Finally, we leverage our framework to re-analyze published scRNA-seq datasets, identifying new regulators that affect the relative abundance of alternatively polyadenylated transcripts, and characterizing extensive cellular heterogeneity in 3’ UTR length amongst antibody-producing cells. Our work highlights the potential for multiplexed single-cell perturbation screens to further our understanding of post-transcriptional regulation *in vitro* and *in vivo*.

## INTRODUCTION

RNA cleavage and polyadenylation represent post-transcriptional regulatory mechanisms that are required for the maturation of eukaryotic pre-mRNA^1–4^. The majority of mammalian genes harbor multiple polyA sites, enabling a single gene to encode multiple mRNA transcripts via alternative polyadenylation^5–7^. The distinct 3’ ends arising from this process add to the rich diversity of mammalian transcriptomes, and can influence multiple distinct stages of the RNA life cycle. For example, shortening of the 3’ untranslated region (UTR) at tandem UTRs can affect transcript stability and localization^8,9^, while alternative polyadenylation at intronic sites can lead to the generation of truncated coding or non-coding transcripts^10–12^. More generally, widespread changes in polyadenylation have been demonstrated in many biological contexts including cellular proliferation^13^, tumorigenesis^14,15^, embryonic development^16^, and secretory cell differentiation^17^.

Biochemical and molecular studies have revealed a subset of core and accessory proteins that are responsible for regulating polyA site choice. For instance, the cleavage and polyadenylation specificity factor complex (CPSF) catalyzes cleavage, the Cleavage factor I (CFIm) and Cleavage factor II (CFIIm) complexes bind auxiliary recognition sequences, and PolyA Polymerase (PAP) is responsible for adding the polyA tail^4^. While the identity of key proteins is known, our understanding of how their relative concentration and interaction with RNA sequence elements influences alternative polyadenylation remains incomplete, representing a key challenge for our understanding of post-transcriptional regulation.

Functional genomics approaches offer exciting potential to address these questions. Recently, massively parallel reporting assays (MPRA) combined with deep neural networks have been utilized to construct sequence-based models of alternative polyadenylation and can successfully predict cleavage site usage under baseline conditions^18–21^. Alternatively, genome-wide 3’ transcriptome technologies can be used to profile changes in polyA site usage across different biological samples^16,22–24^, including those that perform genetic perturbations of CPA regulators. While some individual studies perturb individual or small sets of regulators^25–28^, others have used siRNA-based screening approaches to generate larger resources^29,30^. Alternatively, multiplexed single-cell technologies like Perturb-seq leverage single-cell RNA-sequencing (scRNA-seq) for high-throughput transcriptome-wide characterization of molecular perturbations^31–34^. While scRNA-seq is typically applied to profile heterogeneity in gene expression levels, these data can also be leveraged to characterize changes in transcript structure. In particular, the majority of scRNA-seq protocols are explicitly designed to capture the 3’ end of polyadenylated mRNA transcripts. Therefore, these methods are well-suited to quantify transcriptome-wide polyA site usage at single-cell resolution alongside gene abundances, revealing dynamic changes in polyadenylation during cellular differentiation and disease^35–38^.

Here, we introduced CPA-Perturb-seq, a resource where we perturb known regulators of CPA in a multiplexed 3’ scRNA-seq screen, and quantify each perturbation’s effect on polyA site usage at single-cell resolution. We introduce new computational tools to specifically quantify changes in polyA site usage in sparse single-cell datasets, and to decouple these from changes in gene abundance levels. Our analyses reveal substantial diversity in the number and types of polyA sites affected by perturbation of different regulators, modules of sites that are co-regulated across perturbations, and the role of interacting RNA sequence elements in determining polyA site selection. We also demonstrate how our computational tools can be applied to any 3’ scRNA-seq dataset, identify new regulators in a genome-scale Perturb-seq resource^39^, and characterize natural variation in alternative polyadenylation amongst high-resolution subsets of antibody-secreting plasma cells. Together, our analyses demonstrate how single-cell sequencing can move beyond gene expression analyses and improve our understanding of post-transcriptional gene regulation.

## RESULTS

### Multiplexed Perturb-seq screens of 3’ polyA site usage

We sought to understand how systematic perturbations of genes involved in cleavage and polyadenylation would affect alternative polyadenylation at single-cell resolution (Figure 1A). We designed a library of 162 single guide RNAs (sgRNAs) targeting 42 genes and 10 non-targeting (NT) controls (Supplementary Table 1). Our target set included 18 genes that are known members of core cleavage and polyadenylation complexes, including the Cleavage Factor Im (CFIm), Cleavage Factor IIm (CFIIm), Cleavage and polyadenylation specificity factor (CPSF), and and Cleavage stimulation factor (CSTF) complexes (Figure 1B). We also included 23 genes that have been previously implicated in affecting relative polyA site usage, including subunits of the PAF complex^40^, the splicing factor SRSF3^27^, and THOC5^41^, a member of the transcription/export complex (TREX) (Supplementary Table 1).

**Figure 1:**
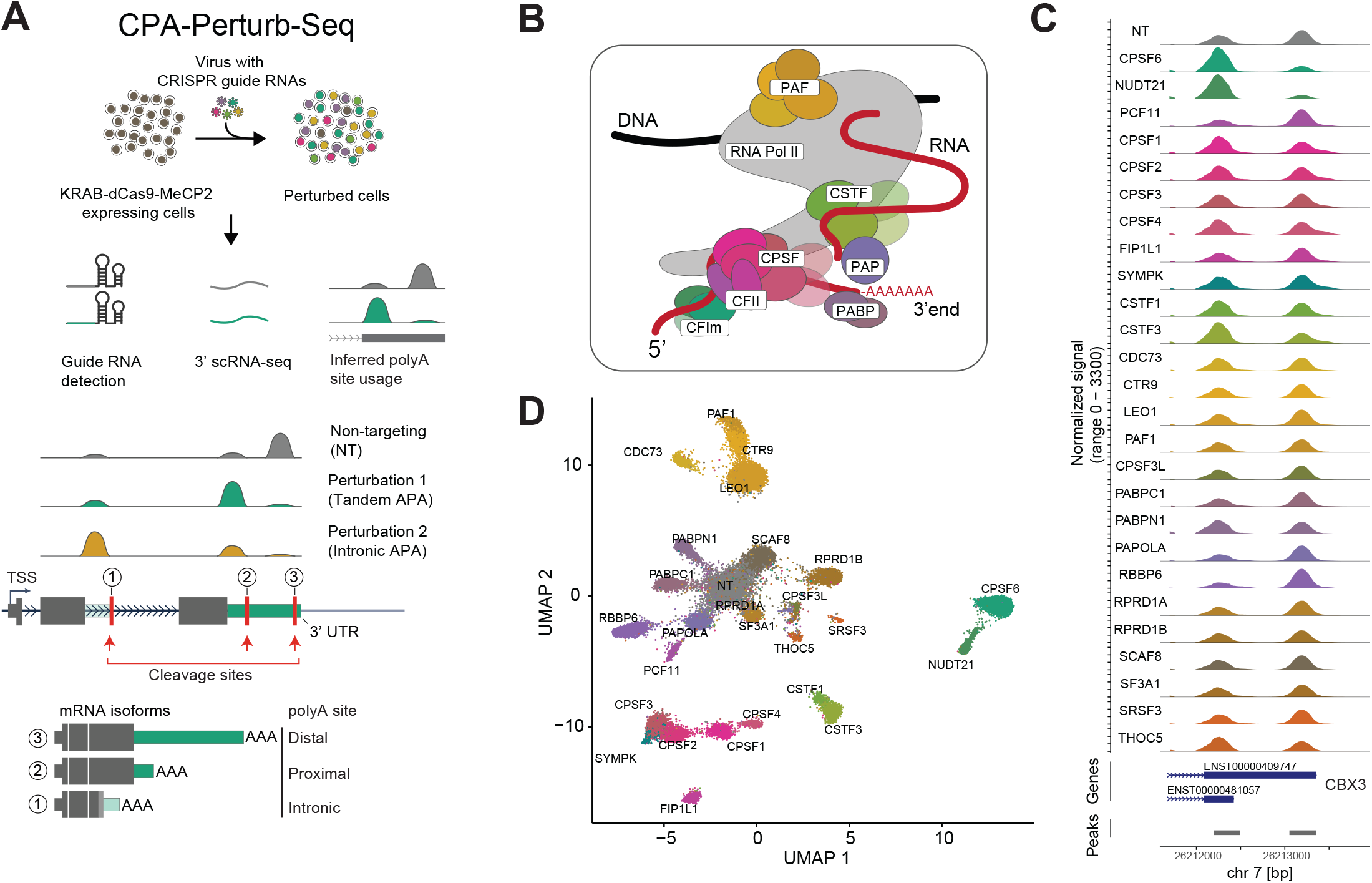
Overview of CPA-Perturb-Seq. **(A)** (Top) Schematic of the experimental workflow used to generate the CPA-Perturb-seq dataset. (Bottom) Schematic of perturbation-dependent changes in either tandem or intronic polyadenylation. **(B)** Diagram depicting core regulatory complexes that make up and interact with the cleavage and polyadenylation machinery. **(C)** Read coverage plots depicting the differential use of alternative polyA sites at the CBX3 locus. Each track represents a pseudobulk average of cells, grouped by their perturbation. ENSEMBL gene models and peaks (quantification region) that precede detected polyA sites are shown below. **(D)** UMAP visualization of HEK293FT cells profiled via CPA-Perturb-Seq. Cells are colored based on the target gene identity. Visualization was computed based on a linear discriminant analysis (LDA) of transcriptome-wide polyA site counts.

We performed a pooled CRISPR inhibition screen (Supplementary Methods) in HEK293FT cells and used the Perturb-seq experimental workflow to simultaneously capture the identity of the guide each cell received along with a 3’ scRNA-seq readout (Figure 1A). While we focus our primary analyses on the deeply profiled HEK293FT dataset (median of 1,788 cells per perturbation), we also repeated the experiment in K562 cells to obtain a second biological context (median of 596 cells per perturbation). Across two biological replicates (independent viral transductions) in each cell line, we obtained a total of 109,661 single cells (Supplementary Table 2) where we were able to successfully assign a single gRNA.

We utilized the scRNA-seq data to quantify both gene expression and transcriptome-wide polyA site usage profiles for each cell. We used tools from the polyApipe pipeline to first identify a set of possible cleavage and polyA sites, and then to quantify their usage in single cells^38^. We further restricted our analysis to polyA sites that are within 50 nucleotides of polyA sites identified in polyAdbv3^7^, a database of polyA sites generated from multiple human cell lines (Supplementary Methods). We only included sites located within an intron or the last exon of a gene (Supplementary Methods). This strategy focuses specifically on splicing-independent changes in alternative polyadenylation, and does not consider changes in alternative last exon usage driven by splicing.

We identified a total of 33,399 polyA sites across 12,194 detected genes, and found that 8,077 genes exhibited usage of two or more polyA sites in our dataset (5,253 genes exhibited usage of three or more) (Supplementary Figure 1A). Moreover, we found that 76% and 79% of our identified exonic and intronic polyA sites, respectively, contained the canonical AATAAA/ATTAAA cleavage motif in the region 50 bp upstream of the inferred cleavage site, as would be expected for *bona fide* polyA sites (Supplementary Figure 1B). We therefore assigned reads from our 3’ scRNA-seq dataset to each polyA site (Supplementary Methods), generating a polyA site/cell count matrix for downstream analysis.

Multiple groups have previously observed that datasets from pooled single-cell CRISPR screens often contain confounding sources of variation^32,33,42^. These include heterogeneity across replicate experiments, cell cycle differences, or variable perturbation efficiency even amongst cells expressing the same gRNA. We applied our previously developed computational pipeline, Mixscape^42^, to address these effects and to remove cells that exhibit no phenotypic evidence of perturbation (Supplementary Methods). For 16 of 42 regulators, Mixscape classified all cells as ‘non-perturbed’, suggesting that even if the perturbation was successful (Supplementary Figure 1C), the global effect on the transcriptome was minimal. For the remaining 26 genes, Mixscape classified 76% of cells as perturbed.

Perturbed cells exhibited diverse changes in alternative polyadenylation. For example, at the CBX3 locus, perturbation of NUDT21 and CPSF6 shifted expression towards the proximal polyA site (‘3’ UTR shortening’) RBBP6 and PCF11 perturbation shifted expression towards the distal isoform (‘3’ UTR lengthening’), while the regulators FIP1L1 and CPSF3L did not induce changes (Figure 1C). These changes were reproducible across biological replicates and multiple independent gRNA (3-4 per gene; Supplementary Figure 1D). We used the polyA site/cell count matrix for perturbed cells along with 3,789 NT controls, as input to linear discriminant analysis (LDA), UMAP visualization (Figure 1D), and unsupervised clustering of the polyA site matrix (Supplementary Figure 1E). These analyses revealed that cells clustered not only by the perturbation they received, but also into broader complexes. For example, cellular profiles after NUDT21 and CPFS6 (both members of the CFIm complex) perturbation were highly correlated, as were profiles for members of the CPSF (CPSF1-4, FIP1L1), CSTF (CSTF1/3), and PAF (PAF1, CTR9, LEO1, CDC73) complexes.

These results suggest our dataset can be used to uncover complex-specific ‘modules’ of co-regulated polyA sites, each of which are responsive to perturbation by a set of functionally related regulators. However, we note that changes in the polyA site/cell count matrix can reflect both changes in 3’ UTR utilization, but also changes in the overall abundance of the gene even in the absence of isoform-level changes. We observed both cases in our dataset. For example, when perturbing CSTF3 (Figure 2A-D), we identify cases where changes in the utilization of a gene’s proximal peak corresponds exclusively to a change in total RNA abundance (ATP6V1G1), exclusively to a change in transcript length due to 3’ UTR shortening (HNRNPH3), or changes in both abundance and relative isoform usage (MRPS16).

**Figure 2:**
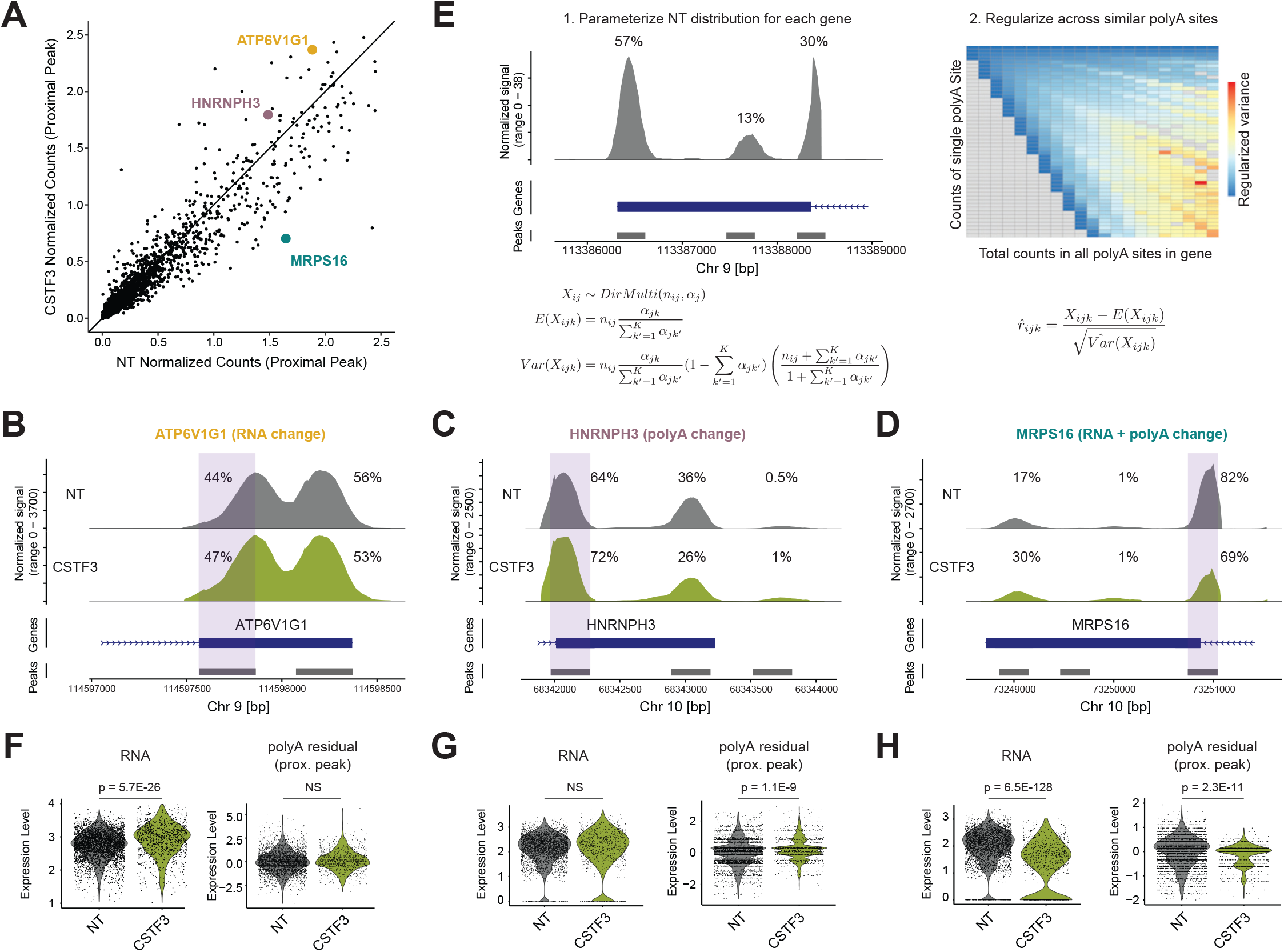
PolyA-residuals quantify alternative polyadenylation at single-cell resolution. **(A)** Average usage of 5,335 proximal polyA sites in NT cells (x-axis) and CSTF3-perturbed cells (y-axis). Only genes with at least two tandem polyA sites are considered. Changes across conditions can breflect eithevr changes in relative polyA site usage, total gene expression, or both. **(B-D):** Read coverage plots at three loci highlighted in (A). Blue box denotes proximal polyA site. **(E)** Schematic depicting the procedure to calculate polyA-residuals (full description in Supplementary Methods). **(F-H)** Violin plots depicting single-cell gene expression levels (left) or single-cell polyA-residuals for the proximal polyA site for NT and CSTF3-perturbed cells. NS (not significant) for RNA comparisons indicate absolute log2FC <0.25 or Bonferonni adjusted p-value >0.05 using Wilcoxon rank-sum test. NS for polyA residual comparisons indicates percent change <0.05 or adjusted p-value >0.05 in differential polyadenylation analysis described in Supplementary Methods.

### Quantifying relative polyadenylation levels at single-cell resolution

To specifically characterize perturbation-driven effects on alternative polyadenylation, we therefore sought to design a computational approach to deconvolve these two effects. While computing ratios of polyA counts for each site within a gene is typically used to study alternative polyadenylation in bulk analyses, computing these ratios in scRNA-seq data is typically infeasible or noisy due to data sparsity. Instead, for each polyA site in each single cell, we aimed to model and quantify the degree of over or under-utilization, compared to the expected usage observed in NT cells.

We note that this problem is conceptually similar to quantifying the degree of increased or decreased expression of each gene in each cell, compared to the population average. We and others have developed statistical methods to address this challenge for gene abundances in scRNA-seq data using generalized linear models^43–46^. Here, we chose to extend this framework to model alternative polyadenylation (Figure 2E). We utilized the Dirichlet multinomial distribution to model the background distribution of each polyA site in NT control cells. The expected value for each site is set by the relative usage of all polyA sites within a gene, which controls for the overall expression of the gene, and can be robustly estimated from 3,789 NT cells.

When quantifying the variance for each site, the Dirichlet multinomial allows for the possibility of overdispersion compared to the standard multinomial^47^, analogous to the routine use of the negative binomial distribution to model Poisson overdispersion when modeling gene abundances^48,49^. This overdispersion accounts for natural biological heterogeneity and ‘intrinsic’ noise that occurs within the background population^46,50^, and can be estimated directly from each dataset. As in sctransform^45^, we first parameterize overdispersion estimates individually for each polyA site, but then regularize these estimates across similar sites (Supplementary Methods). The output of our procedure is a statistical model for each polyA site, describing its background usage across NT control cells.

Finally, we utilized these background models to quantify relative polyA site usage at single-cell resolution. By comparing the observed counts at each site in each cell with the expected value and variance from the Dirichlet multinomial model, we compute a Pearson residual (‘polyA-residual’) at each polyA site. The sign and magnitude of this residual describes the cell’s relative deviation from the expected background distribution for each polyA site. A positive residual reflects that a polyA site is used more frequently in a single cell relative to the background distribution, and a negative residual reflects that a polyA site is used less frequently than the background distribution.

Our quantified polyA-residuals can be used as input for differential polyadenylation analysis, allowing us to identify polyA sites whose relative frequency changes across groups of cells while mitigating any confounding changes in overall gene abundance. We tested for changes in polyA-residuals using a linear model, including gRNA identity as a covariate, to identify reproducible changes for each perturbation (Supplementary Methods). When applied to our previous examples (Figure 2F-H), this approach successfully distinguished loci where we observed either changes in transcript structure, abundance, or both. The polyA-residuals can also be used as input for clustering and visualization. When repeating our LDA-based visualization procedure and clustering analyses on the polyA-residual matrix (Supplementary Figure 3A), we replicated our previously observed findings (Figure 1D) confirming that these co-regulatory patterns were driven by coordinated changes in polyadenylation.

**Figure 3:**
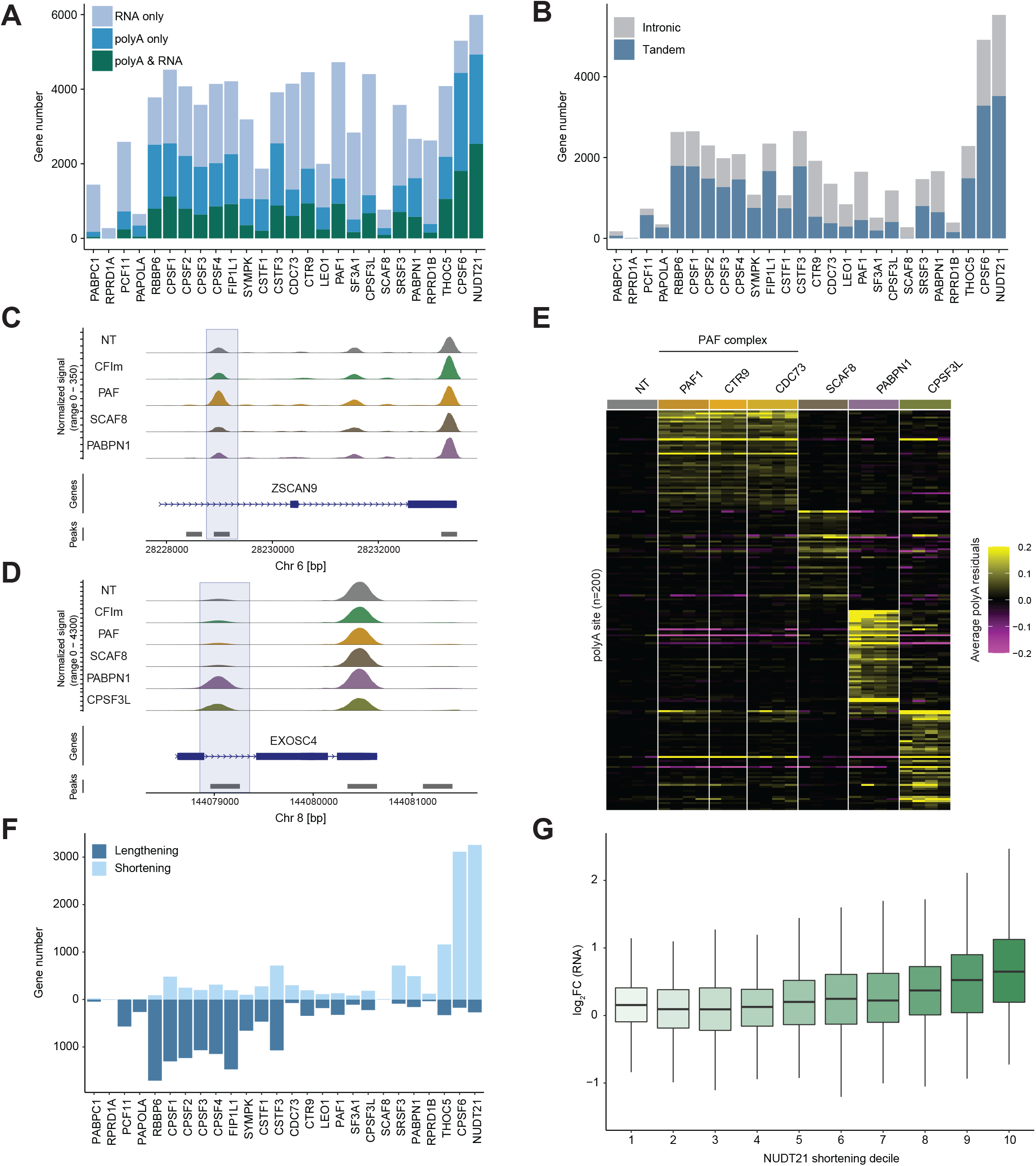
Characterizing tandem and intronic alternative polyadenylation in CPA-Perturb-Seq. **(A)** Number of genes with significant changes in RNA abundance, relative usage of at least one polyA site, after perturbation of each regulator. Barplots show results in HEK293FT cells. **(B)** Total number of genes with relative changes in intronic or tandem polyA site usage in HEK293FT cell dataset. **(C-D)** Read coverage plots showing differential usage of intronic sites (boxed) at the ZSCA9 (C) and EXOSC4 locus (D). **(E)** Heatmap showing polyA residuals for intronic sites that are uniquely differentially utilized after perturbation of PAF, SCAF8, PABPN1, and CPSF3L. Each heatmap cell shows the pseudobulk average of cells after grouping by sgRNA identity. **(F)** Number of genes with significant changes in tandem polyA site usage in HEK293FT cell dataset, classified by 3’ UTR shortening or 3’ UTR lengthening. **(G)** Boxplot indicating the observed log2 fold-change in gene expression after NUDT21 perturbation. Genes are partitioned into deciles based on the degree of 3’ UTR changes observed after NUDT21 perturbation.

### Characterizing perturbation-dependent changes in polyadenylation

We next characterized the effect of each perturbation individually. We identified 6,734 genes that exhibited differential alternative polyadenylation (at least one polyA site with differential usage) in at least one the 26 gene perturbations, but observed substantial differences across regulators. CFIm complex members such as NUDT21 exhibited the strongest perturbation responses affecting more than 5,600 genes (Figure 3A), including 2,397 genes where we exclusively detected relative changes in at least one polyA site (40%; blue bar), 1,058 genes where we exclusively detected changes in total transcript levels (18%; light blue bar), and 2,536 genes where we detected changes in both abundance and structure (42%, green bar).

Our results demonstrated that differential analysis of our polyA-residuals represents an effective workflow to identify specific changes in relative polyA site usage, as opposed to total transcript abundance. For example, we observed that perturbation of PABPC1, affected the expression level of 1,265 genes, but had negligible effects on relative polyA site usage in either our HEK293FT or K562 cell dataset (Figure 3A; Supplementary Figure 3C). This result is consistent with the known cytoplasmic localization of PABPC1, which binds the polyA tail after nuclear export, and is not expected to regulate polyA site choice^51^.

We next classified each significant change in polyA site usage as reflecting either intronic polyadenylation or tandem polyadenylation, based on site annotations in the polyADB database (Figure 3B). We found that in most cases, perturbing an individual regulator primarily led to changes in tandem polyA site usage. However, for a minority of regulators, such as polymerase-associated factor (PAF) complex members^52^ or the RNA PolII elongation factor SCAF8^26^, responses were primarily associated with intronic polyadenylation in both HEK293FT and K562 cells (Supplementary Figure 3D). As both sets of regulators interact with RNA Polymerase and play an established role in regulating polymerase progression, these results provide additional evidence for kinetic models where changes in elongation rate can influence alternative polyadenylation^53^, and suggest that this relationship is particularly important in the context of intronic sites.

While we identified multiple regulators that primarily affected intronic polyadenylation, we found that they regulated distinct intronic sites. (Figure 3C-E). Moreover, we found that the distance between two adjacent cleavage sites in a transcript was predictive of the responsiveness to PAF1 perturbation (Supplementary Figure 3E). This relationship was strongest for polyA sites located in the first intron, but also held for downstream sites as well (Supplementary Figure 3E). However, when performing similar analyses for SCAF8, we observed a weaker predictive power for both intronic location or distance between cleavage sites (Supplementary Figure 3F). These results demonstrate that while transcriptional elongation rate likely influences intronic polyA site selection, polymerase-interacting factors can exhibit distinct regulatory effects.

Focusing next on alternative polyadenylation between tandem polyA sites, we found that perturbation of CPA regulators resulted in striking shifts in the utility of either proximal or distal polyA sites, with most perturbations (18/26) exhibiting a skew of greater than 70% in either direction (Figure 3F). These relationships replicated in our independently obtained K562 dataset as well (Supplementary Figure 3G). We did observe a general trend where 3’ UTR shortening was associated with an increase in total gene abundance (Figure 3G), consistent with the broad association between 3’ UTR length and the presence of regulatory elements that may impact RNA stability^54,55^.

Our observed patterns of shortening/lengthening were concordant with previous studies that utilize bulk 3’ end sequencing technologies, but highlighted the advantages of the Perturb-seq technology. For example, four previous studies^25,29,30,56^ have consistently revealed that perturbation of NUDT21 perturbation affects polyA site usage in a subset of genes (ranging from 375-1,600) and leads to 3’ UTR shortening at tandem UTRs. In our dataset (Figure 3F; Supplementary Figure 3G), we observed more than 5,500 genes exhibiting significant changes in polyA site usage after perturbation, exhibiting not only high sensitivity but also high specificity in both HEK and K562 datasets (>93% of tandem UTR changes resulted in shortening, indicating that these reflect *bona fide* perturbation responses).

Similarly, RBBP6 perturbation has been associated with 3’ UTR lengthening, but the degree of this preference (ranging from 60% to 78%) and the number of genes (ranging from 100 to 1,300) varies across studies^28–30^. In our dataset, we observed more than 2,600 genes with polyA site changes, with a high specificity (>95% lengthening at 3’ UTR) in both cell lines (Figure 3F; Supplementary Figure G). These results highlight how pooled single-cell CRISPR screens, which avoid batch effects by multiplexing all perturbations and controls together, can yield accurate perturbation signatures especially when performed with high cell number and utilizing multiple independent gRNA. Moreover, this experimental design is ideally suited for the identification of co-regulated polyA sites across multiple regulators, without having to compare datasets generated across different experiments or studies.

### Modules of co-regulated polyA sites exhibit distinct functional properties

While our previous analyses characterized regulators individually, we also clustered perturbations based on the observed changes to all differentially polyadenylated sites quantified by our model (Supplementary Methods; Figure 4A). Consistent with expression-based analysis, perturbation clusters reflected membership structure of core CPA complexes, as well as additional evidence of co-regulation. For instance, RBBP6, FIP1L1, and PCF11 are not members of the same complex, but all cause 3’ UTR lengthening at overlapping sites upon perturbation, and cluster together. Moreover, we repeated these analyses on the K562 dataset and observed highly concordant correlation patterns (Figure 4B), suggesting these reflect co-regulatory relationships that generalize beyond a single biological context.

**Figure 4:**
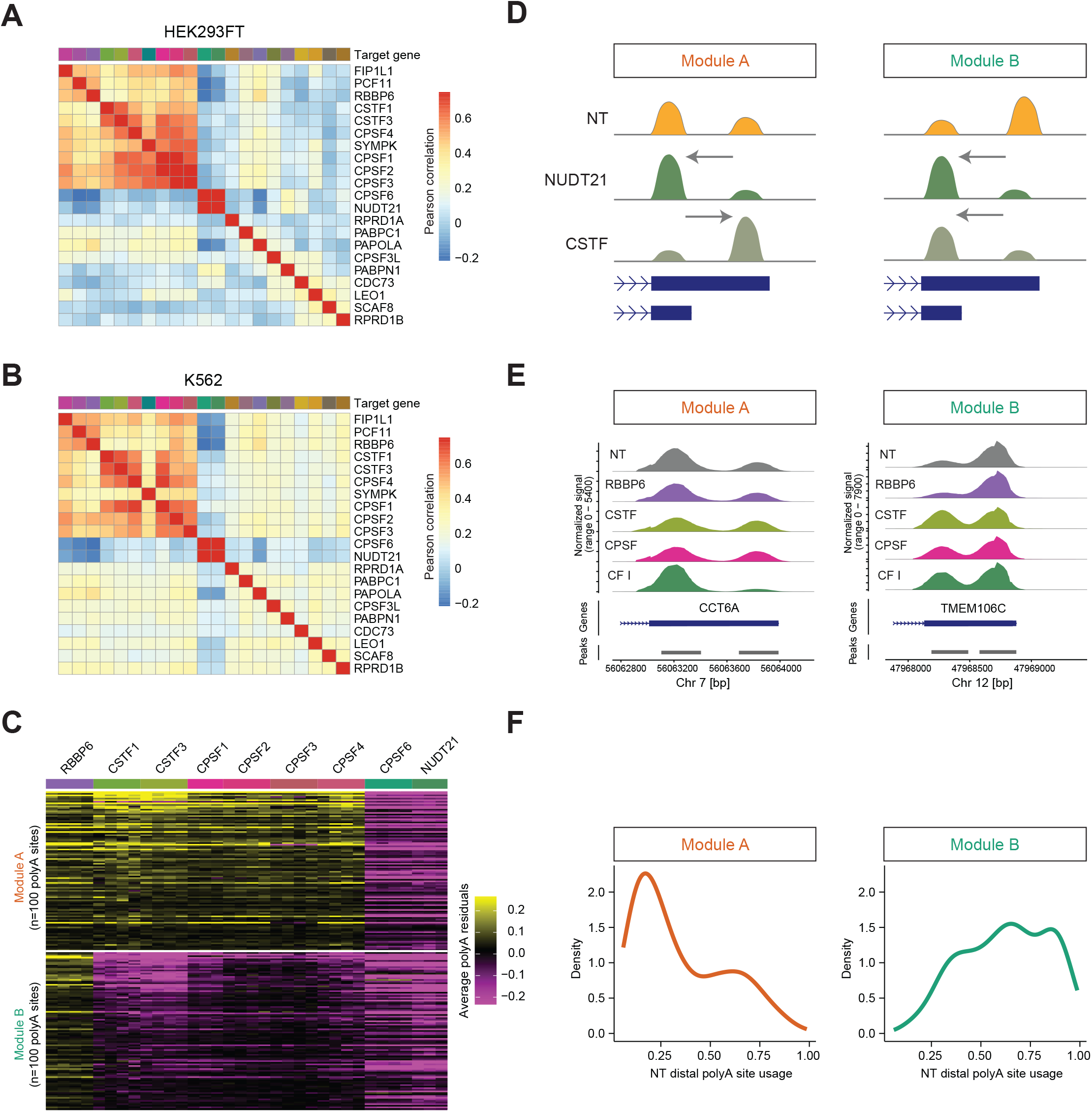
Modules of co-regulated polyA sites exhibit functional differences. **(A)** Pearson correlation matrix depicting the relationships between perturbations in HEK293FT cells. Correlations are calculated using the linear model coefficients learned during differential polyadenylation analysis (Supplementary Methods). Matrices include all perturbations where we obtained at least 50 cells in both HEK and K562 cells, and are ordered via hierarchical clustering. **(B)** Same as (A), but the correlation matrix is generated from an independent analysis on K562 polyA residuals. **(C)** Heatmap showing polyA-residuals for distal peak sites in Module A genes (CSTF and CPSF act in the opposite direction as CPSF6/NUDT21), and Module B genes (CSTF and CPSF act in the same direction as CPSF6/NUDT21). For visualization, the top 100 polyA sites, ranked by the magnitude of CSTF perturbation, are shown for each module. **(D)** Schematic diagram of genes belonging to module A and Module B. **(E)** Read coverage plots showing polyA site usage of representative genes belonging to module A (left, CCT6A) and module B (right, TMEM106C). **(F)** Density plot showing distal site usage in NT control cells for genes belonging to Module A (left) versus Module B (right). Genes in Module A tend to use the proximal site, while genes in Module B tend to use the distal site.

We found that the correlation structure was not exclusively driven by global preferences towards shortening and lengthening, but also local differences in the specific sites affected by each regulator. For example, perturbation of RBBP6 (preference towards 3’ UTR lengthening) and CFIm complex members CPFS6 and NUDT21 (preference towards 3’ UTR shortening) showed strongly anti-correlated responses, reflecting their globally opposing regulatory tendencies at the same set of loci. By contrast, CSTF and CPSF complex members (preference towards 3’ UTR lengthening) showed only weak anti-correlation with CFIm members, reflecting more complex patterns of co-regulation.

To further explore this, we considered a group of 1,208 genes that exhibited transcriptional shortening after CFIm perturbation (Supplementary Methods). When further subdividing this set of sites based on their response to CSTF perturbation, we observed an expected module (Figure 4C-E, Module A) of 245 polyA sites (20%) where CSTF perturbation resulted in an opposing lengthening response. However, we also identified a module (Module B) of 110 (9%) genes where CSTF perturbation also resulted in shortening, phenocopying CFIm perturbation despite their opposing global preferences. The remaining 71% of sites did not exhibit changes in utilization upon CSTF perturbation. While we identified these modules in our HEK293FT dataset, we independently observed reproducible patterns at the same loci in K562 cells (Supplementary Figure 4A).

Strikingly, we found that these gene modules exhibited clear functional differences (Figure 4F). In particular, we found that genes where we observed opposing regulatory effects between the two complexes (Module A) strongly favored the usage of proximal sites in NT cells, while genes exhibiting consistent regulatory effects (Module B) were strongly biased towards distal site usage. These results were consistent in both HEK293FT and K562 cells (Supplementary Figure 4B) and indicate that local effects, likely determined by differences in sequence content, establish the responsiveness to CSTF perturbation, and are important in establishing the proximal versus distal bias for individual genes. More broadly, we conclude that our polyA residuals represent an effective statistical approach for characterizing the perturbation responses of individual regulators, and for identifying modules of polyA sites that are co-regulated across perturbations.

### APARENT-Perturb reveals an interactive cis-regulatory code

Our identification of modular patterns of differential polyadenylation that reproduce across cell types emphasizes the role of local sequence drivers in determining an individual polyA site’s responsiveness to distinct perturbations. Motivated by the success of deep learning models in accurately predicting genome-wide patterns of alternative polyadenylation in baseline conditions^18–20,57–59^, we sought to extend these models to predict the perturbation responses observed in our dataset. For example, APARENT2 represents a residual neural net, originally trained on MPRA datasets, that can predict baseline polyA site usage in HEK293FT cells and interpret specific sequences elements and genetic variants that drive model accuracy^57^. The ability to successfully capture nonlinear interactions, including positional and combinatorial interdependencies between motifs, highlights the ability of these models to learn intricate cis-regulatory determinants^60^.

We therefore hypothesized that sequence-based learning models could predict the response of each polyA site to each of the ten highest magnitude perturbations in our dataset. To test this, we first used the pre-trained APARENT2 model to provide baseline predictions for polyA site usage. We then trained a new neural network (APARENT-Perturb) to predict usage in our Perturb-seq data, using 200nt sequences centered on the site of 3’ cleavage, along with the baseline APARENT predictions as input (Figure 5A). This approach was inspired by the MTSplice model^61^, and represents an ensemble-based multi-task perturbation network that can not only predict relative polyA site usage in NT control cells (baseline), but also can predict polyA site usage after perturbation. After training, APARENT-Perturb could accurately predict the isoform proportion of polyA sites for held-out genes in both the non-targeting (NT) condition (R_S_ = 0.70) and in perturbations (0.65 ≤ R_S_ ≤ 0.73 depending on perturbation), as measured by 10-fold cross-validation (Figure 5B, Supplementary Figure 5A). When predicting relative differences in polyA site usage between a given perturbation and the NT condition, the performance varied more as some perturbations resulted in an only moderate change to APA levels (0.27 ≤ R_S_ ≤ 0.59), but these results were still highly significant (2.25×10^−125^ ≤ p ≤ 2.66×10^−17^).

**Figure 5.**
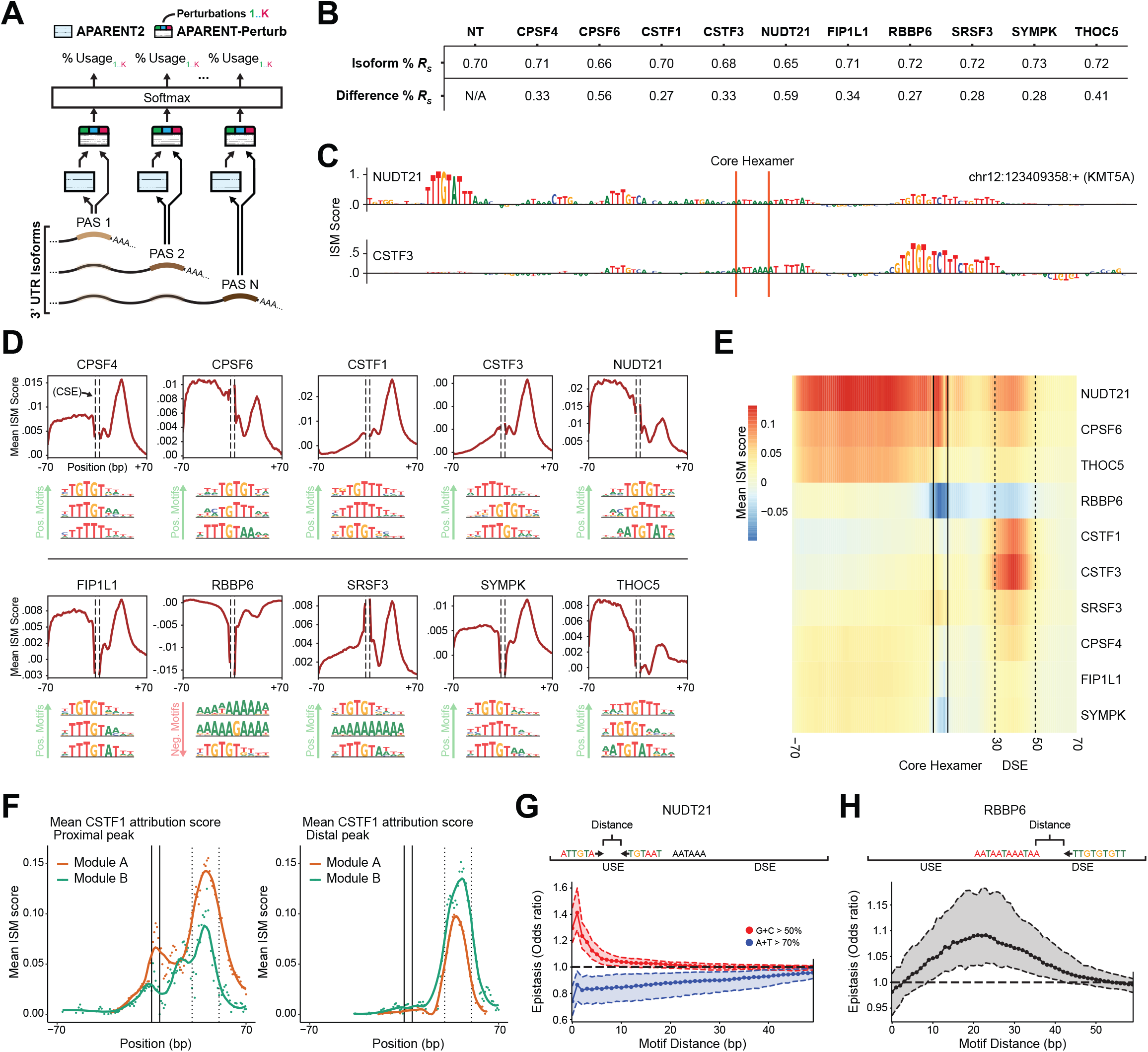
A multi-task neural network predicts perturbation responses from RNA sequence. **(A)** Schematic of (APARENT-Perturb), an ensemble-based neural network architecture for predicting perturbation responses. Green/blue/red output heads correspond to model predictions for the K perturbation conditions. **(B)** 10-fold cross-validation performance when predicting distal isoform proportions (top row) or differences in distal isoform proportion with respect to the NT condition (bottom row). **(C)** Sequence-specific attribution scores for two example perturbations in the KMT5A gene. Attribution scores are displayed after calculating residuals with respect to NT cells. **(D)** Averaged attribution scores as a function of position, for 10 perturbations. The three top MoDISco motifs are shown for each perturbation (Supplementary Methods). **(E)** Heatmap showing averaged attribution scores for each perturbation, as a function of position, for the distal-most site in each gene. **(F)** Mean attribution scores for CSTF1 perturbation in Module A vs Module B for both proximal sites (left) and distal sites (right). Location of the core hexamer and downstream sequence elements (DSE) are marked with solid and dashed vertical lines, respectively. Plots show the mean attribution score at single base-pair resolution (points), as well as the loess-smoothed trend (lines). **(G)** Epistasis analysis for dual UGUA motifs, in either G/C-rich contexts (red) or A/U-rich contexts (blue). The y-axis reflects the effect on predicted NUDT21 perturbation after dual insertion of both motifs, compared to the effect of inserting one motif at a time (Supplementary Methods). **(H)** Epistasis analysis of canonical hexamers and U/G-rich motifs, based on the RBBP6 perturbation.

To interpret the model, we performed *in silico* mutagenesis (ISM) by simulating local sequence alterations and comparing the resulting predictions to the unaltered model. This procedure yields a set of nucleotide-level ‘attribution scores’, reflecting the contribution of each individual base to the model’s prediction^62,63^. Importantly, by subtracting scores of the NT (baseline) output, we isolate each sequence’s importance in predicting perturbation responses. For example, the attribution scores of the distal polyA site in the KMT5A gene, highlight an upstream UGUA motif that is predicted to drive responsiveness to NUDT21 perturbation, and a distinct downstream GU-rich region motif that drives responsiveness to CSTF3 perturbation (Figure 5C, Supplementary Figure 5B). For each perturbation, we averaged ISM scores across loci to identify regions that harbored important sequence elements (Figure 5D-E). We next used a motif discovery tool, TF-MoDISco^60,64^, to cluster the attribution scores of each perturbation into a set of salient motifs (Figure 5D, Supplementary Figure 5C-D). These results recapitulate and extend previously established binding motifs and positions^2,4,59^, for example, NUDT21 and CPSF6 both display high average importance in the upstream region of polyA sites and are sensitive to UGUA motifs with T- or A-rich flanks, while CSTF1 and CSTF3 display a peak of importance in the downstream region with U- or GU-rich sequences among their top motifs.

Intriguingly, APARENT-Perturb attribution scores suggest motifs that help to coordinate joint activities of both CFIm and CSTF complex members. Our model’s attribution scores predicted that CFIm perturbation responses are predicted not only by sequences upstream of the cleavage sequence, but also by a sequence element located approximately 30-50 bp downstream (downstream element; DSE). This DSE overlaps with a region of predicted importance for CSTF perturbation, reflecting a co-enrichment of functional sequences for both complexes at the same sites (Supplementary Figure 5E). While APARENT-Perturb predicted positive attribution scores for most sites in this region after NUDT21 perturbation (Supplementary Figure 5F; Decile 10), a subset (Decile 1) exhibited negative attribution scores. Indeed, we found that these two groups of sites differed in their responsiveness to CSTF1 and CSTF3 (Supplementary Figure 5G). The fact that the NUDT21 perturbation model ascribes importance to sequence motifs that drive CSTF regulation is strong evidence of sequence-driven interaction between these factors.

We had previously observed that CSTF and CFIm complex members can jointly regulate polyA sites in either the same or opposing directions (Figure 4C-F), and we identified a link between our model’s attribution scores and our previously identified modules. Specifically, we found that in genes where CFIm perturbation led to transcriptional shortening and CSTF perturbation led to lengthening (Module A), the DSE at the proximal polyA site was characterized by sequence elements with high CSTF attribution scores, which are predicted to facilitate CSTF binding and regulation. However, at genes where perturbation of both complexes led to transcriptional shortening (Module B), the proximal sites exhibited significantly weaker sequence elements (Figure 5F left, p< 2.0*10^−5^, Wilcoxon two-sided rank sum test). By contrast, we observed increased attribution scores for Module A genes at distal sites (Figure 5F right, p< 1.6*10^−4^). Taken together, these findings suggest a model where the sequence content at proximal polyA sites is particularly important both in establishment of the proximal/distal bias, as well as the responsiveness to multiple perturbations. In a subset of genes (Module A), proximal peaks contain strong motifs that serve to recruit CSTF. This regulatory structure promotes cleavage at the proximal site under baseline conditions, but leads to transcriptional lengthening after CSTF perturbation. Alternatively, a distinct gene subset (Module B) exhibits weaker sequence features at the proximal site, while the distal site contains sequences that promote recruitment of CSTF and CFIm. These loci exhibit distal cleavage under baseline conditions, but perturbation of either complex results in transcriptional shortening.

Finally, as neural networks trained on ChIP-seq data have been recently shown to successfully learn the syntax of cooperative binding between transcription factors^60^, we aimed to identify similar types of interactions between CPA regulators. We used APARENT-Perturb to simulate either individual or pairwise motif insertions, and compared the predicted results to identify epistatic interactions. For example, the CFIm complex includes a NUDT21 homodimer, but it is unclear if and how multiple UGUA motifs affect binding^65,66^. We found that two adjacent UGUA motifs tended to act cooperatively in predicting the responsivity to NUDT21 perturbation. However, we only observed synergistic effects when both motifs were surrounded by GC-rich sequences, while an AT-rich context was associated with sub-additive interactions (Figure 5G, Supplementary Figure S5H-I). We also identified that two distinct sequence elements, the canonical core hexamer, and GU-rich sequence element observed in the DSE, also exhibited epistasis in predicting polyA site usage after RBBP6 perturbation. While previous work has associated that both of these motifs independently are associated with RBBP6 regulation^28^, APARENT-Perturb identified a position-dependent relationship, with a maximum epistatic interaction observed when the motif distance was approximately 20bp (Figure 5H, Supplementary Figure 5J). We verified each of these results using polynomial feature regression (Supplementary Figure 5K-L). We conclude that deep learning models can be successfully applied to analyze high-throughput Perturb-seq datasets, and can reveal a cis-regulatory landscape that encodes complex patterns of co-regulation across multiple complexes.

### Identification of APA regulators from genome-wide screening datasets

While the genes selected for our screen encompassed previously identified regulators of alternative polyadenylation, our computational workflow is capable of characterizing changes in polyA site usage for any 3’ scRNA-seq dataset. We therefore reanalyzed a recently published genome-wide Perturb-seq dataset (GWPS)^39^ which perturbed 9,866 transcriptionally active genes in K562 cells (including all 26 perturbed regulators in our datasets), but did not explore perturbation-dependent changes in polyA site usage. To address this, we computed polyA-residuals for each cell, and used these as input to differential polyadenylation analysis (Supplementary Methods). As the GWPS dataset contained far fewer cells per perturbation (median 91 cells, vs. 1,032 for in our dataset for the 26 overlapping perturbations), we identified substantially fewer genes exhibiting changes in polyA site usage (median of 165 genes per overlapping perturbation, compared to 1,351 in our data). However, even at shallow depth, the GWPS dataset enabled accurate global characterization of each regulator. For example, we observed a strong concordance in the global bias towards 3’ UTR shortening or lengthening induced by regulatory perturbation across both datasets (Figure 6A).

**Figure 6:**
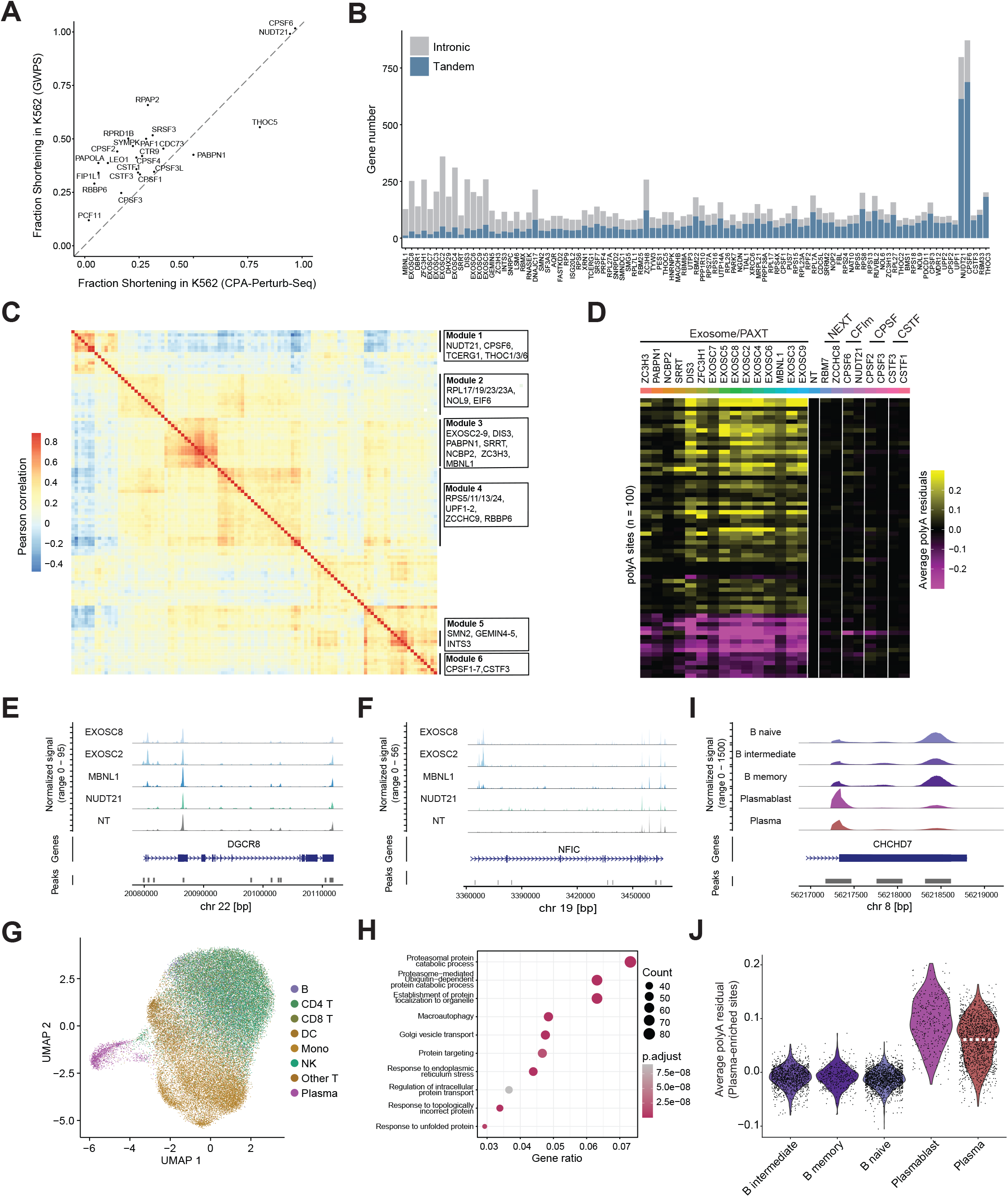
Characterizing heterogeneity in relative polyA site usage in additional 3’ scRNA-seq datasets. **(A)** 3’ UTR shortening preference observed after perturbing regulators in the CPA-Perturb-seq dataset (x-axis), and the GWPS dataset (y-axis). We observe concordant results for this global metric across datasets. **(B)** Same as Figure 3B, but for the GWPS dataset. **(C)** Correlation matrix depicting the relationship between perturbations in the GWPS dataset, as in Figure 4A. Representative genes for each of the six correlated modules are shown on the left. All genes are listed in Supplementary Figure 6A. Shown are all perturbations where we detected changes in relative polyA site usage in at least 50 genes. **(D-F)** MBNL1 perturbation phenocopies perturbation of PAXT complex members. **(D)** Heatmap shows polyA-residuals for polyA sites that are differentially utilized after both MBNL1 and PAXT perturbation. **(E-F)** Representative read coverage plot depicting changes in polyA site usage after perturbation of PAXT complex members and MBNL1. **(G)** UMAP visualization generated from polyA residuals of PBMC dataset. Cells are colored based on their gene expression-based cell annotation. **(H)** Gene ontology enrichment analysis on genes exhibiting 3’UTR shortening in plasma cells compared to B cells. **(I)** Read coverage plot depicting 3’UTR shortening in CHCHD7 gene in distinct B and plasma cell subpopulations. **(J)** Average polyA-residual (reflects degree of 3’ UTR shortening) of proximal sites with increased usage in plasma cells. We observe extensive heterogeneity within the plasma cell lineage, including increased shortening in cycling plasmablasts, and two subpopulations of non-cycling plasma cells (denoted by horizontal line, Supplementary Figure 7C).

We therefore extended our analyses to focus on a previously annotated set of 1,280 RNA binding proteins^67^ in order to facilitate identification of regulators that directly modify RNA. While most perturbations exhibited minimal transcriptome-wide changes in alternative polyadenylation, we identified 172 regulators whose perturbation affected polyA site usage in at least 50 genes (Figure 6B, Supplementary Table 4). We also identified groups of highly correlated perturbations that were consistent with and substantially expanded our previous observations (Figure 6C, Supplementary Figure 6A). For example, one group included the CFIm complex members CPSF6 and NUDT21, but also THOC3, a Transcription-Export (TREX) complex member, and TCERG1 (transcriptional elongation regulator 1). While perturbation of this module was associated with shortening at tandem 3’ UTR (Supplementary Figure 6B-C), we identified a separate lengthening-associated module (Module 4; Supplementary Figure 6F) consisting of Up-frameshift complex members (UPF1, UPF2), the small ribosomal subunit (RPS24, RPS4X), and the ribosome maturation factor TSR2. While these genes are well-studied regulators of translational control and RNA stability, none have been previously associated with regulating polyA site selection. Components of the large ribosomal subunit formed were also associated with polyA site selection, but formed a separate module (Module 2; Supplementary Figure 6D), along with the translation initiation factor EIF6. These analyses demonstrate how large-scale perturbation screens can identify novel regulatory factors and suggest tight regulatory crosstalk linking changes in alternative polyadenylation with multiple processes in the RNA life cycle.

We additionally identified a third group of 15 correlated perturbations (Module 3), 13 of which have been previously identified as members of the poly(A) tail exosome targeting (PAXT) complex^68^. Perturbation of this module was associated primarily with the indirect up-regulation of intronically polyadenylated transcripts (Figure 6B), whose abundance was not changed in response to perturbation of the nuclear exosome targeting (NEXT) complex or the CPA machinery (Figure 6D). This response is driven by the PAXT complex’s role in degrading prematurely terminated RNA transcripts, which accumulate in the cytoplasm after PAXT perturbation^68,69^, although the nuclear surveillance machinery that specifically distinguishes premature transcripts remains unknown^70^. Intriguingly, the nuclear cap-binding complex member NCBP2 and and splicing regulator MBNL1 were also members of this module but neither are members of the PAXT complex. While NCBP2 is known to promote successful RNA export^68,71^, MBNL1 perturbation has been previously linked to regulating levels of intronic retention^72^, including in cases where retention leads to premature termination^73^. The striking phenocopying between perturbation of MBNL1 and PAXT subunits (Figure 6D-F) suggests a hypothesis where a splicing regulator, via its role in regulating intron retention, may assist PAXT in selecting unstable and undesired transcripts for degradation.

### scRNA-seq profiles reveal extensive plasma cell heterogeneity in polyA site usage

While our previous analyses focused on cellular heterogeneity across *in-vitro* perturbation experiments, we next asked whether our statistical framework could quantify and interpret APA heterogeneity in an *in-vivo* context. For example, recent work using bulk RNA-seq datasets has demonstrated that secretory cell differentiation is associated with widespread changes in polyadenylation^17^. To further explore this, we calculated polyA-residuals on a 3’ scRNA-seq dataset consisting of 49,958 circulating human peripheral blood mononuclear cells (PBMC) which includes seven COVID-19 infected samples that exhibit an induction of antibody-secreting plasma cells^74^. We processed this data to identify and quantify 20,067 polyA sites, and quantified both gene expression levels as well as polyA residuals.

Unsupervised analysis of polyA residuals revealed that heterogeneity in polyA site usage across immune subsets was relatively modest compared to heterogeneity in gene expression (Figure 6G). However, consistent with previous reports^11,17^, we did observe that plasma cells exhibited clear differences compared to all cell types, including developmentally related B cell subsets. In addition to observing a shift towards the shorter isoform of the IGHM locus (one of the first described examples of alternative polyadenylation^75^; Supplementary Figure 7A), we identified 1,8783 genes (630 intronic changes and 1253 tandem changes) exhibiting differential usage of at least one polyA site in plasma cells (Supplementary Methods). Genes exhibiting differential tandem polyadenylation were primarily associated with 3’ UTR shortening (95%). Gene Ontology analysis revealed a strong enrichment for Golgi vesicle transport and protein localization, which are linked to the core secretory phenotypes of plasma cells (Figure 6H).

One key advantage of single-cell measurements is the ability to explore how multiple levels of granularity in cell annotation affect downstream analyses. We found that when further subdividing plasma cells to annotate short-lived and highly proliferative plasmablast subpopulations, these cells exhibited the most striking shifts in polyA site usage (Figure 6I-J, Supplementary Figure 7B). Surprisingly, we found that non-cycling plasma cells not only exhibited weaker changes, but also exhibited a bimodal distribution in their polyA-residuals, enabling further subdivision into two groups based on the degree of 3’ UTR shortening (Figure 6J; Supplementary Figure 7C-D). These two groups differed not only in transcript structure, but also in the expression of a module of genes that were highly enriched for their involvement in respiratory and metabolic processes (Supplementary Figure 7E). These results extended previous bulk RNA-seq based findings^17^, but were uniformly consistent across 11 donors (Supplementary Figure 7F). They demonstrate that widespread 3’ UTR remodeling occurs in the earliest stages of plasma cell differentiation, but substantial cellular heterogeneity in polyA site usage remains even after commitment to this lineage. Future experiments will establish whether the metabolic changes observed between these groups relate to the secretory capabilities or lifespan of these cells.

We conclude that 3’ scRNA-seq data can be combined with tailored computational pipelines to explore cellular heterogeneity in polyA site usage for both in vitro and primary samples, and have developed an open-source R package PASTA (PolyA Site analysis using relative Transcript Abundance) that implements the analytical methods described in this manuscript. PASTA is fully compatible with our analytical toolkit Seurat^76^, and the software release includes a vignette demonstrating how users can explore changes in their datasets using PASTA and Seurat. These data and code resources will facilitate the characterization of heterogeneous alternative polyadenylation in diverse biological systems and a deeper understanding of the sequences and regulatory factors that govern post-transcriptional regulation.

## DISCUSSION

In this study, we aimed to understand how the abundance of CPA regulators, as well as the presence of RNA sequence elements, affect the regulation of alternative polyadenylation across the transcriptome. We demonstrate that the Perturb-seq technology, which has been widely utilized to study transcriptional regulatory networks, can be successfully applied to study post-transcriptional regulation as well. We introduce a statistical framework to quantify changes in relative polyA site usage at single-cell resolution, and demonstrate how this approach can characterize the effect of individual regulators, identify modules of co-regulated polyA sites, and enumerate subpopulations of cells that exhibit changes in polyA site usage in any 3’ scRNA-seq dataset.

Our CPA-Perturb-seq dataset revealed striking heterogeneity in the perturbation responses of different regulators. This was reflected in the number, type, and directionality of changes induced by each perturbation. However, our dataset highlights that regulation of alternative polyadenylation is not a uniform or global process, where all polyA sites are sensitive to perturbation by all core regulators. Instead, we consistently observed evidence for modularity and substructure in our data. Even when perturbing regulators of the core CPA machinery, we identified groups of polyA sites that were co-regulated by a subset, but not all, regulators. Moreover, we identified cases where the same set of perturbations resulted in opposing responses for distinct modules of polyA sites. Our Perturb-seq dataset is well-suited for module characterization, as the multiplexed design mitigates experimental batch effects and avoids the need to compare perturbation profiles generated from different experiments or studies.

By interpreting our multi task deep neural network, APARENT-Perturb, we find that this local regulatory structure is encoded in part by sequence-specific elements that surround the cleavage site. Previous models trained on MPRA data constitute a powerful approach to identify functionally important sequence elements, but it is challenging to understand how they exert regulatory effects. By integrating these models with our perturbation data, we learn direct associations between sequence elements and regulators, providing a more mechanistic understanding of cis-regulatory element function. We demonstrate the ability of this approach to identify interactions between different regulators of alternative polyadenylation, but this approach could also be extended to deep neural networks that predict chromatin accessibility levels from DNA sequence, and to provide deeper functional interpretation of sequence variants.

While our analyses aimed to focus on regulatory mechanisms that influenced cleavage and polyadenylation decisions, we repeatedly observed cases where additional regulatory processes in the RNA life cycle would alter the relative abundance of alternatively polyadenylated transcripts. For example, we observed regulator-specific patterns that connected changes in RNA polymerase elongation rate with altered usage of intronic polyA sites. More broadly, we also found that perturbation of proteins with well-characterized roles in RNA export, RNA translation, and RNA splicing and intron retention also resulted in differential usage of polyA sites. These results highlight the extensive interdependencies that connect different RNA regulatory processes. To this end, future work may be able to exploit these interdependencies to infer RNA kinetic parameters from 3’ scRNA-seq data, for example, utilizing changes in the usage of intronic polyA sites to infer cell type-specific changes in RNA elongation rate. More broadly, our statistical method may be extended to characterize additional sources of transcriptomic diversity, such as changes in splicing from full-length datasets, in order to characterize a broader realm of post-transcriptional regulatory events.

While our study independently explores datasets deriving from either multiplexed perturbation screens or primary human samples, looking forward, we believe these contexts will be mutually informative. Functional genomics tools like Perturb-seq are especially well-suited to identify causal relationships between molecular regulators and their targets. In contrast, comparative analysis of alternative polyadenylation across biological samples, conditions, and disease states is a powerful approach for identifying transcriptome-wide changes, but identifying the causal regulators driving these responses remains challenging. We envision that the molecular signatures inferred from experiments where causal relationships are established represent important resources to interpret molecular signatures where causal relationships are unknown. These links will be particularly informative as Perturb-seq experiments extend beyond *in-vitro* models, as we perform here, towards true *in-vivo* settings. Integration of these datasets therefore represents a potential path forward for systematic reconstruction of the regulatory factors and networks that govern post-transcriptional regulation and the RNA life cycle.

## DATA AND CODE AVAILABILITY

The CPA-Perturb-seq datasets generated for this manuscript are available for download at: https://zenodo.org/record/7619593#.Y-P7Zi1h2X0

Seurat and PASTA are both available as open-source R packages at: https://github.com/satijalab/seurat

https://github.com/satijalab/PASTA

Code to train and interpret the APARENT-Perturb model is available at https://github.com/johli/aparent-perturb

## ACKNOWLEDGEMENTS

The authors would like to acknowledge Torben Heick Jensen, Robert Bradley, Christina Leslie, and Christine Mayr for thoughtful discussions related to this work. We thank the Neville Sanjana lab for providing access to the KRAB-dCas9-MeCP2 plasmid. The work was supported by the Chan Zuckerberg Initiative (EOSS5-0000000381, HCA-A-1704-01895 to R.S.), and the NIH (RM1HG011014-02, 1OT2OD033760-01 to R.S). A.K. and J.L. were supported by NIH grants 2U24HG007234.

