## Supplementary Methods for "CPA-Perturb-seq: Multiplexed single-cell characterization of alternative polyadenylation regulators"

#### Cell culture and maintenance

HEK293FT (HEK) and K562 cells were acquired from ThermoFisher (R70007) and ATCC (CCL-243), respectively. Monoclonal HEK293FT and K562 lines that constitutively express KRAB-dCas9-MeCP2 (CRISPRi) were generated as previously described[1,2]. HEK and K562 cells were maintained in DMEM (Caisson DML23) or RPMI (Thermo Fisher 11875119), respectively, supplemented with 10% fetal bovine serum (Serum Plus II Sigma-Aldrich 14009C) and with no antibiotics at 37°C and 5% $CO_2$. Constitutive CRISPR Cas effector expressing cells were maintained using 5μg/mL Blasticidin S (ThermoFisher A1113903). Guide RNA expressing cells were grown using 1μg/mL puromycin (ThermoFisher A1113803).

#### sgRNA design, virus production and virus transduction

We selected the most effective sgRNAs from previous genome-wide CRISPR-interference screens[3,4] along with non-targeting sgRNAs that showed no depletion. Individual sgRNAs were cloned into lentiGuideFB-Puro-A (Addgene #192506) as previously described[2]. All constructs were confirmed by Sanger sequencing. Individual plasmids were pooled accounting for the relative expected essentiality of target genes to compensate for the expected loss of cells with essential gene sgRNAs.

For pooled virus production we seeded $1x10^7$ HEK293FT cells per 10 cm dish 12-18 hours before transfection [60μl PEI, 6.4 μg psPAX2 (Addgene #12260), 4.4 μg pMD2.G (Addgene #12259) and 9.2 μg of the plasmid pool]. Six to eight hours post-transfection, the medium was exchanged for 10 ml of DMEM + 10% FBS containing 1% bovine serum albumin (BSA). Viral supernatants were collected after additional 48 hours, spun down to remove cellular debris for 5 min at 4°C and 1000 x g and passed through a 0.45 μm filter prior to storage at -80°C.

For single-cell experiments we transduced $3x10^6$ HEK293FT or K562 CRISPRi cells per 12-well with an MOI of  < 0.1 (< 10% survival) ensuring high coverage (cells per sgRNA) of at least 1000x and a single integration probability > 95%. The cells were selected with 1μg/ml puromycin starting at 24 hours post-transduction for at least 48 hours for HEK293FT cells and at least 5 days for K562 cells. Cells were maintained with blasticidin and puromycin until the single-cell experiment 6-7 post transduction.

#### Direct capture Perturb-seq and sequencing

We performed scRNA-seq (10x Genomics Chromium Single Cell 3' Gene Expression v3 with Feature Barcoding technology for CRISPR screening) 6-7 days post-transduction. Before the run, cell viability was determined (>95%). We leveraged Cell Hashing in order to super-load (~40,000 cells/lane) the Chromium instrument[5]. Gene expression and sgRNA feature libraries were constructed following the manufacturer's protocol. Cell Hashing libraries (Hashtag-derived oligos, HTOs) libraries were prepared following the recommended protocol on cite-seq.com. All libraries were sequenced on an Illumina NovaSeq S4 flow cells.

### Processing of single cell gene expression data

HEK and K562 scRNA-seq and Feature Barcoding data was processed using CellRanger (v3.0.1, ‘cellranger count’). HTO data was quantified using the *CITE-seq-count* package (v1.4.2). Count matrices were then used as input into the Seurat R package (v4.0) to perform downstream analyses. We retained cells where we detected between 2,500 to 8,000 (HEK) and 1,000 to 7,000 genes (K562), and where less than 15% of UMI originated from mitochondrial genes. HTO and sgRNA counts were normalized using the centered log-ratio transformation approach. We retained cells with unique HTO and sgRNA assignments generated using the *MULTIseqDemux*[6] function in Seurat.

### Quantifying polyA site usage

CellRanger output BAM files were used as input to polyApipe (https://github.com/MonashBioinformaticsPlatform/polyApipe), a computational workflow which quantifies polyA site usage at single-cell resolution. Briefly, this workflow comprises two steps.

First, polyApipe attempts to discover a set of polyA sites that are utilized in the dataset. All reads in the overall sample that contain a stretch of at least 5 softclipped A’s at the 3’ end of the read that were not present in the genomic sequence are annotated as ‘polyA reads’. This subset of reads is used for peak calling. We required at least 5 ‘polyA reads’ across all cells to form a polyA site. In cases where two independently discovered sites fell within the defined counting region (here: 300 nucleotides upstream of the identified polyA site) the peak with the higher polyA read count was retained.

Second, polyApipe quantifies the read coverage, at each of these polyA quantification windows , for each single cell in the dataset. For counting, all reads were used, regardless of whether they were originally annotated as ‘polyA reads’. The outputs of polyApipe are a tabular count file and counts matrix quantifying the usage of each polyA site at single-cell resolution, as well as a GFF file that describes the location of the inferred polyA sites. These outputs can be read via the PASTA package using the function *ReadPolyAPipe*.

To further ensure that our downstream analysis was conducted on *bona fide* polyA sites, we intersected these sites with those in PolyA_DB version 3.2 (polyAdbv3)[7,8], a catalog of sites identified from deep sequencing data using the (3'READS) method. We lifted over these sites from hg19 to hg38. We retained all polyA sites identified by polyApipe that were located within 50 nucleotides of a polyAdbv3 polyA site. We additionally used polyAdbv3 to assign a gene annotation to each polyA site, and further restricted our downstream analyses to polyA sites that are either located within an intron, or the last exon, of a transcript. The same set of polyA sites was used for analysis in K562 cells by providing the HEK GFF file as an input to polyApipe.

In order to visualize the usage of polyA sites, we used the blocks function in Sinto (https://github.com/timoast/sinto) to construct a fragment file from the aligned BAM file. The function *CoveragePlot* in Signac was used to visualize read coverage throughout this manuscript (i.e. Figure 1C). In the bottom of these plots, we visualize the peak locations that represent the polyA sites identified by polyApipe.

### Mixscape analysis and visualization

We have previously developed a computational toolkit, Mixscape[9], to analyze and interpret Perturb-seq datasets. Mixscape aims to address confounding sources of heterogeneity in these datasets, and to identify cells that may have received a sgRNA but do not exhibit strong molecular evidence of successful perturbation. We use the polyA counts matrix as input to Mixscape analysis. Briefly, for each regulator, Mixscape considers all cells receiving a sgRNA targeting that regulator, and models them as a Gaussian mixture model with two components. The parameters of the first component are constrained to reflect the distribution of NT control cells. The second learned component should therefore reflect successfully perturbed cells. For each cell that receives a gRNA, Mixscape calculates a posterior probability for each component. Cells that were classified (posterior probability >0.5) as belonging to the first component are similar to NT control cells ('non-perturbed'), and therefore discarded from further analysis.

In order to visualize our dataset, we used the MixscapeLDA function. All perturbed and NT cells were used as input for this analysis. Linear discriminant analysis (LDA) is a supervised dimensional reduction technique that attempts to find a low-dimensional subspace that maximally discriminates between different groups ('perturbations') in our data. As previously described[9], to prevent overfitting during this procedure, we reduce the dimensionality of our dataset to 130 features prior to running LDA (five projected principal components * 26 perturbations). Twenty-six components returned from LDA are used as input for 2D visualization in UMAP (Figure 1D).

### A single-cell metric for polyA site usage (polyA residuals)

We sought to develop a single-cell metric for polyA site usage that adjusts for total abundance of a gene in order to prioritize the identification of relative changes in polyA site usage. We focus our analyses on genes that contain more than one polyA site, and leverage NT cells in order to estimate a background distribution describing the usage (counts) of each polyA site. We utilize the Dirichlet multinomial distribution, allowing us to model overdispersion (relative to the multinomial distribution) that frequently arises in the context of scRNA-seq, due to both biological and technical heterogeneity. The parameters of the Dirichlet multinomial include both the total number of reads observed across all polyA sites, as well as a vector $\alpha$ that yields a probability distribution parameterizing the relative usage of different polyA sites.

Our approach consists of three steps, as described in detail below. (1) Fitting Dirichlet multinomial parameters for background polyA site usage within each gene individually, using NT cells (2). Regularizing variance estimates across similar polyA sites (3). Comparing polyA sites observed in each perturbed single cell to this model and calculating model residuals (polyA residuals).

*<u>Fitting Dirichlet multinomial parameters for each gene individually</u>*

We define the following:

$X_{ijk}$: number of counts at polyA site $k$ ($1 \leq k \leq K$) for gene $j$ in cell $i$

$n_{ij}$: total number of reads in cell $i$ across all polyA sites in gene $j$: $n_{ij} = \sum_{k=1}^{K} X_{ijk}$

$\alpha_j \in \mathbb{R}^K$: vector of Dirichlet parameters for each of *K* polyA sites, where each $\alpha_{jk} > 0$

Then, for each cell $i$, we model the counts for all polyA sites within gene $j$ using the Dirichlet Multinomial distribution.

$$X_{ij} \sim DirMulti(n_{ij}, \alpha_j)$$

For each gene, we independently estimate the Dirichlet parameters, $\hat{\alpha}_j$, using count data from the 3,789 NT cells. We obtained parameter estimates using the MGLMfit function in the MGLM package[10] (dist = "DM"), which obtains maximum likelihood parameter estimates using iteratively reweighted Poisson regression[11].

Once we have obtained the parameter estimates , $\hat{\alpha}_j$, we can calculate the expected counts falling into each polyA site, as well as the variance of these counts under our background model:

$$E(X_{ijk}) = n_{ij} \frac{\alpha_{jk}}{\sum_{k'=1}^{K} \alpha_{jk'}} \quad \text{Eq. 1}$$

$$Var(X_{ijk}) = n_{ij} \frac{\alpha_{jk}}{\sum_{k'=1}^{K} \alpha_{jk'}} (1 - \sum_{k'=1}^{K} \alpha_{jk'}) \left( \frac{n_{ij} + \sum_{k'=1}^{K} \alpha_{jk'}}{1 + \sum_{k'=1}^{K} \alpha_{jk'}} \right) \quad \text{Eq. 2}$$

We calculate these values for each polyA site in each NT cell.

*Regularizing variance estimates in NT cells across similar polyA sites*

As has been repeatedly been observed for both bulk and scRNA-seq data, individual variance estimates obtained from generalized linear models with overdispersed error distributions can be noisy and subject to overfitting[12–14]. Therefore, after obtaining estimated variances for our background model, we regularize these estimates across similar polyA sites in order to increase robustness.

We reasoned that polyA sites could be considered similar not only if they were utilized at a similar relative abundance within a gene, but also if the overall gene abundance was similar. We therefore regularize across two dimensions, the expected value of the number of reads falling into each polyA site, $E(X_{ijk})$, as well as the total number of reads that fall into polyA sites in that gene, $n_{ij}$.

To regularize, we first define a 30 x 30 non-overlapping grid with evenly spaced intervals from the minimum of each dimension to the 99th percentile in NT cells. Each bin is defined by a range of evenly spaced parameter values (intervals) for each of the two dimensions. After defining this grid, we perform the following procedure for each NT cell: we assign each of the polyA sites to one of the bins based on two dimensions, $E(X_{ijk})$ and $n_{ij}$. For each polyA site, we add estimated variance from Eq. 2 to the bin. After completing this procedure for all NT cells, we sample 1000 observations (polyA sites) from each bin if the number of observations in that bin exceeds 1000. Then, we use a multivariate kernel regression estimator, as implemented in the pcf.kernesti function in the regpro package (http://jklm.fi/regpro/). This calculates a smoothed variance, at the midpoint of each point of the 30x30 grid. We use this midpoint as the regularized variance ($\hat{Var}(X_{ijk})$) for each polyA site falling within that bin.

<u>*Calculating polyA residuals for all cells*</u>

Given our background model estimated from NT control cells, we can now ask the following questions for any single cell $i$ in our dataset: given the observed read depth $n_{ij}$ observed for this cell at gene $j$:

1) Under the background model, what is the expected value of the counts at each polyA site
2) Under the background model, what is the variance for these counts
3) How do the observed values for this cell compare with the expected values?

The expected value for each site can be obtained from Eq. 1 above. The variance under the background can be obtained by identifying the correct bin for each polyA site (based on the two dimensions $E(X_{ijk})$ and $n_{ij}$, and obtaining the regularized variance estimate for that bin. If the dimensions for the polyA site fall outside of the grid defined in the previous step (defined by the 99th percentile in NT cells), we use Eq. 2 to calculate the variance. We also set a minimum regularized variance of 0.1 for all polyA sites.

To compare the observed counts from each cell to the expected values under the background model, we calculate a Pearson residual (polyA residual), $\hat{r}_{ijk}$, representing a standardized distance between the observed and expected counts.

$$\hat{r}_{ijk} = \frac{X_{ijk} - E(X_{ijk})}{\sqrt{\hat{Var}(X_{ijk})}}$$

We calculated two sets of Pearson residuals: one based on tandem sites located in the 3' most exon, and one on all intronic sites and tandem sites in the 3' most exon. This allowed for the identification of both tandem and intronic alternative polyadenylation events.

By calculating residuals at the polyA site level (instead at the gene level), our procedure can flexibly handle diverse cases, for example, when there are >2 polyA sites in a single 3' UTR, or when a gene exhibits differential polyadenylation at both intronic and tandem polyA sites. However, we note that the polyA residuals at multiple sites in the same gene are not independent (i.e. if one polyA site exhibits high relative usage and receives a positive residual within a single cell, other polyA sites will likely decrease in relative use and receive negative residuals). For this reason, when we perform downstream analyses such as clustering or principal components analysis (which often assume independence across features) using polyA residuals, we only consider a single polyA site per gene.

**Model-based tests for differential polyA site usage**

In order to identify polyA sites whose relative usage changes after each perturbation, we performed differential analysis on the polyA residuals using a linear model. For each site $k$ and each perturbation, we ran a linear model comparing the perturbed cells to NT cells with main effects for each of $l$ guides that target the same gene.

$$\hat{r}_{ijk} = \beta_0 + \beta_1 Z_{i1} + \ldots + \beta_l Z_{il},$$
$$Z_{il} = 1 \text{ if cell } i \text{ received guide } l, \, 0 \text{ otherwise}$$
$$Z_{i1} = \ldots = Z_{il} = 0, \text{ when cell } i \text{ received NT guide.}$$

For each site, we only include cells with non-zero values for the residuals in the linear model, representing cells in which at least one read aligned to polyA sites in the same gene was captured. We then prepare a contrast in order to compare all of the guides targeting the same gene to NT cells, as well as comparisons between guides. For instance, if there are 3 guides that target a particular gene, the contrast matrix will be as follows:

$$\begin{pmatrix} -1 & 0 & 0 \\ \frac{1}{3} & 1 & 0 \\ \frac{1}{3} & -1 & 1 \\ \frac{1}{3} & 0 & -1 \end{pmatrix}$$

The first column compares NT cells to the average across each guide. The second column compares the first and second guides, and the third column compares the second and third guides. By weighting each guide equally we will downweight observations where the perturbation strength is not observable across multiple independent gRNA. In order to test the specified contrasts matrix, we use it as input to the contrasts argument to the lm function in R. The corresponding estimate ('perturbation effect estimate') and p-value for the contrast comparing NT cells to all guides is then used to assess if a particular site responds to a perturbation.

Linear modeling was performed separately on both the tandem sites and intronic polyA residuals. After running this linear model on all polyA sites, the q-value package (https://github.com/StoreyLab/qvalue) was used to estimate q-values with a false-discovery rate (FDR) of 5%. Genes having at least one tandem site with a q-value less than 0.05 and greater than 5% change in average usage compared to NT cells were classified as tandem alternative polyadenylation events. Similarly, genes with at least one intronic site with a q-value less than 0.05 and at least 5% change compared to NT cells were classified as intronic differential polyadenylation events. For Figure 3B, a gene could have both intronic and tandem differential polyadenylation events.

We also utilize the results of the linear model to perform clustering on the alternative polyadenylation perturbation responses across regulators. To do this, we first select all genes that show evidence of alternative polyadenylation in any perturbation, and extract the top polyA site (based on the magnitude of the polyA residual) for each gene. For each regulator, we define a vector of perturbation effect estimates (obtained from our linear model contrasts) for each of these polyA sites. These vectors were used as input to hierarchical clustering in Figures 4A-B.

In order to intersect changes in polyA site usage with potential changes in gene abundance, we also estimated total RNA levels for each gene with multiple polyA sites. We summed counts across all annotated polyA sites for that gene for each cell, regardless of position within the transcript, and performed standard log-normalization in Seurat. We then performed differential expression to compare gene expression of each perturbation to NT cells, using the Wilcoxon Rank sum test. Significance was determined using a false discovery threshold of 0.05, as above. Genes were defined as differentially expressed for a perturbation if the log2FC exceeded 0.25 and the q-value was less than 0.05.

**Analysis of intronic polyadenylation sites**

When performing downstream analysis of perturbation-induced changes in intronic polyadenylation, we focused on members of the PAF complex, SCAF8, PABPN1, and CPSF3L. For each perturbation, we identified intronic sites that had an increase in average usage >5% compared to NT cells as well as q-value <0.05, as obtained in our linear modeling framework above. We then identified sites that were significant for only one intronic regulator, and plot the top 50 unique intronic sites for each regulator in Figure 3E.

For the PAF complex, we sought to assess if the strength of response of each polyA site to PAF perturbation was related to the distance to the next polyA site. For each intronic polyA site, we calculated the distance to the next annotated polyA site in the gene. We then demonstrated that distance to the next polyA site predicted PAF1 response by binning polyA sites into deciles based on the distance to the next polyA site. We then calculated the Pearson correlation between the distance (log-scale) and the percent change of the intronic polyA site in PAF1 perturbed cells compared to NT cells (Supplementary Figure 3E). We also performed this analysis separately for polyA sites located within first introns, and all other introns, after obtaining first intron annotations from the knownCanonical table obtained from the UCSC genome browser (hg38). We also repeated the same analysis for SCAF8 (Supplementary Figure 3F).

**Defining Modules Based on CSTF/CPSF responsiveness**

We aimed to first identify a set of sites that were responsive to NUDT21 perturbation (shortening), but could in principle result in lengthening in response to other perturbations. We identified 1208 polyA sites located in the last exon whose usage was decreased in response to NUDT21 perturbation and increased in response to RBBP6 perturbation, using a threshold of q <0.05 and a percent change compared to NT that was greater than 1%. These sites represent predominantly distal sites who usage decreased in CF-I perturbation and increased in RBBP6 perturbation.

We then sought to classify these sites on the basis of how they responded to perturbation of additional core complexes, in particular, CSTF and CPSF. To do this, we ran an additional linear model in order to identify a robust set of peaks that respond similarly across members of the same complex. We ran the model as described above, testing for the difference between NT cells and guides targeting a gene, but grouped together all members of the same complex (CPSF1-4 and FIP1L1 for CPSF and CSTF1/3 for CSTF).

Then, we identified a set of sites that were upregulated in both CSTF and CPSF perturbations (using the complex-based linear model described above), representing 245 genes. These genes compose Module A, where CPSF/CSTF perturbation acts in the opposite direction of NUDT21 perturbation. We also identified 110 polyA sites composing Module B, whose usage decreased in CSTF and CPSF perturbation, indicating that NUDT21 perturbation and CPSF/CSTF perturbation induced 3' UTR shortening at these genes.

**APARENT-Perturb architecture, training and interpretation**

*Data processing*

The perturbation data was aggregated to pseudo-bulk level by grouping cells by knock-down gene. The polyA sites of each gene were intersected against the coordinates of annotated sites in PolyADB V3[15]. Only sites within the 3' UTR were retained. Sites without a well-defined core hexamer motif (at most two substitutions away from the canonical AAUAAA motif) were removed. Finally, genes with less than two retained sites, or more than 10 sites, were thrown out. The resulting filtered data consisted of 5,267 genes. For each gene and perturbation, we used the isoform read counts to estimate the relative usage (proportion) of each polyA site.

*Architecture and training*

The neural network follows a residual architecture with 6 layers of dilated convolutions (32 filters per layer, each filter is 3 positions wide), taking as input 205 bp of sequence in a one-hot-encoded format centered on the 3' cleavage site and outputs residual logit scores for the NT condition and the 10 chosen perturbations. The architecture differs slightly from the tissue-specific multi-PAS model proposed in APARENT2[16]. Specifically, the neural network does not only output one logit score per sequence and perturbation, but instead it outputs three scores per perturbation: a proximal site score, a middle-site score and a distal site score. This allows the model to learn different cis-regulatory rules of polyA site usage depending on site type. The model is replicated and shared across all polyA sites of a given gene (for a maximum of 10 sites), such that the model predicts a tensor of perturbation response logits $S \in \mathbb{R}^{10 \times 10 \times 3}$ given an input tensor of polyA site sequences $X \in \{0, 1\}^{10 \times 205 \times 4}$. The response logits S are used in combination with baseline APARENT2 logits $A \in \mathbb{R}^{10}$ and the log distance between sites, $D \in \mathbb{R}^{10}$, to predict the final proportion of usage for each polyA site and perturbation, $\hat{Y} \in [0, 1]^{10 \times 10}$:

$$\hat{y}_{i,j} = \frac{\mathbb{1}_{\{\text{PAS } i \text{ exists}\}} \times \exp\Big(\text{score}(i, j)\Big)}{\sum_{t=1}^{10} \mathbb{1}_{\{\text{PAS } t \text{ exists}\}} \times \exp\Big(\text{score}(t, j)\Big)}$$

Where:

$$\text{score}_{i,j} = \left[\sum_{k \in \{\text{p,m,d}\}} \mathbb{1}_{\{\text{PAS } i \text{ is type } k\}} \times w_k^{(\text{score})} \times \big(a_i + s_{i,j,k}\big)\right] + w^{(\text{distance})} \times d_i + w_{i,j}^{(\text{bias})}$$

And:

$i$: Site # in 3' UTR
$j$: Perturbation #
$k$: Proximal, middle or distal output
$a_i$: APARENT2 score for site $i$
$d_i$: Cumulative log-distance (bp) from proximal-most site
$s_{i,j,k}$: Predicted score of perturbation $j$ at site $i$ for type $k$ (trainable)
$\boldsymbol{w}$: Regression weights (trainable)

Similar to the training procedure described in APARENT2, we first fit the regression weights **w** while keeping the perturbation-specific scores $s_{i, j, k}$ locked at zero. During this phase, we minimize the KL-divergence between measured proportions Y and predicted proportions Y^. After convergence, we unfreeze the scores $s_{i, j, k}$ and fit the weights of the underlying neural network using a combination of a KL-divergence loss and a margin error between measured differences ($Y_{Perturb}$ - $Y_{NT}$) and predicted differences ($Y\hat{}_{Perturb}$ - $Y_{NT}$). The model was trained with Keras using the Adam optimizer[17] and training stopped once the loss started to increase on held-out validation data.

*Windowed In-silico Saturation Mutagenesis and motif clustering*

Attribution scores were generated for the polyA sites of all 5,267 genes and for each perturbation using a windowed In-silico Saturation Mutagenesis (ISM) approach[18,19]. By sliding a 5 nt wide window over each sequence, we randomly and uniformly mutate all 5 bases within the window and record the mean difference in predicted logit scores with respect to the wildtype sequence for 8 independent samples. The resulting mean log odds ratio is taken as the importance score for the base at the center of the window. Whenever the importance scores for a particular perturbation are analyzed, we always subtract the importance scores of the non-targeting (NT) condition from the scores of the perturbation.

To cluster the importance scores and discover motifs, we ran TF-MoDISco[20] on the scores of each perturbation with the following settings: sliding window = 8, flank size = 5, max seqlets = 40,000, FDR = 0.1 and # mismatches = 1). TF-MoDISco was executed separately on the negative- and positive part of the importance scores.

*Motif insertion analysis, pairwise ablations and polynomial feature regression*

Dual motif insertion simulations were performed on a set of 64 wildtype polyA site sequences as backgrounds. These sites were uniformly randomly sampled from all distal sites unless otherwise specified (e.g. in some analyses we biased the selection to GC- or AT-rich contexts). For each wildtype sequence, we predicted the residual score $r^{(WT)} = s_{Perturb} - s_{NT}$ for the perturbation of interest. We then inserted motif A, motif B or motif A and B at every possible combination of positions (*i* and *j*) and generated the corresponding predictions $r^{(A)}_{i, j}$, $r^{(B)}_{i, j}$, $r^{(A\text{ and }B)}_{i, j}$. Given these quantities, we estimated the odds ratio of epistasis at positions *i* and *j* as: $epi = \exp((r^{(A\text{ and }B)}_{i, j} - r^{(WT)}) - ((r^{(A)}_{i, j} - r^{(WT)}) + (r^{(B)}_{i, j} - r^{(WT)})))$. Finally, by grouping pairs of positions by their distance (| i - j |), we estimated the 10th, 50th and 90th percentile of odds ratios for each distance.

Pairwise motif ablations were estimated using an identical formula, except now motifs are ablated (mutated) from wildtype sequences where they already exist. Motifs were ablated by replacing the seqlet identified by MoDISco with uniformly random bases for 32 independent samples, unless the motif occurred over highly conserved positions (e.g. the core hexamer), in which case the motif was randomly exchanged for specific hand-crafted subsequences (e.g. weaker version of the hexamer motif).

Polynomial feature regression was used to validate the epistatic relationships discovered by the neural network directly in the data. To validate the epistasis of a given pair of motifs A and B, we constructed independent count features of the occurrence counts of motif A and motif B respectively, as well as binary indicator features of higher-order combinatorial co-occurrences (e.g. both motif A and B occur n times in site *i*). These features were constructed for each polyA site in each gene. To reduce the number of free parameters, the features for all non-distal sites in the same gene were aggregated. Given this feature matrix as input, we performed standard linear regression in scikit-learn[21] to regress the log of the odds ratio of distal polyA site usage in the perturbation of interest relative to the NT condition. To interpret the learned epistatic rules, we inspect the signs and magnitudes of both the individual count feature weights and the combinatorial binary indicator weights. For example, to assess the homotypic relationship between instances of motif A, we perform regression using a count feature of motif A and a binary indicator which is set to 1 only if two or more instances of motif A are present in the sequence. After regression, if the learned weight for the binary indicator is negative it suggests a sub-additive relationship (the regression subtracts from the additive count feature). Conversely, a positive indicator weight suggests a super-additive relationship (the regression adds to the additive count features).

**Analysis of Genome-scale Perturb-seq dataset**

We processed the scRNA-seq gene expression raw data from the Genome-scale Perturb-seq dataset[22] (K562 KD8) using CellRanger (v6.0.0, 'cellranger count'; hg38, ensembl v97). Guide RNA and target gene assignments were extracted from the metadata of the original analysis (https://gwps.wi.mit.edu). We quantified polyA site counts as described previously (*Quantifying polyA site usage),* using the same polyA site GFF file as in the HEK293FT dataset. From this dataset, we extracted cells with gRNA targeting 1,280 previously annotated RNA binding proteins[23], along with NT controls. We kept all polyAsites located within 50bp of a previously annotated site in either the polyAdbv3[15] or polyAsite 2.0[24] database. After intersection, we retained sites whose annotated location was either within an intron or the last exon.

We performed differential polyadenylation for each perturbation as previously described for our CPA-Perturb-seq dataset (*Model-based tests for differential polyA site usage*). After linear modeling, we defined differential APA events based on a FDR-threshold of 0.05. For all perturbations with more than 50 APA events, we calculate a correlation matrix using the same procedure used in Figure 4A-B. In order to focus on perturbations that cluster together, we retain regulators that exhibit a correlation of >0.25 with at least one other perturbation, and visualize this correlation matrix in Figure 6B.

**Analysis of human peripheral blood mononuclear cell dataset**

We obtained scRNA-seq reads from GSE155673[25] , and performed alignment and gene expression quantification using CellRanger (v6.0.0, 'cellranger count'; hg38, ensembl v97). We retained all cells with at least 200 detected genes. Cells were annotated on the basis of their RNA profiles using the Azimuth reference for PBMC, available at (https://azimuth.hubmapconsortium.org/references/). Mapping was performed independently for each donor. We excluded cells annotated as platelets and erythroid cells, resulting in 49,958 cells across 11 donors. We identified polyA sites in this dataset using the same procedure as we followed for our CPA-Perturb-seq dataset (*Quantifying polyA site usage*).

We computed polyA residuals for each cell as described above, using all cells to estimate a background distribution. To obtain the UMAP visualization in Figure 6G, we follow a procedure similar to unsupervised scRNA-seq analysis. First, we estimate a set of 'variable' features. To do this, we calculate the variance (excluding zero values) of polyA residuals across cells for each polyA site. We then rank polyA sites by their variance. We use this ranking to select 2,000 variable sites, but select at most one polyA site from an individual gene to ensure that each feature remains independent. After variable feature selection, we perform scaling, centering, and clipping as in scRNA-seq, followed by principal components analysis. The first 30 PCs were used as input to UMAP visualization in Figure 6G.

In order to identify polyA sites that drive plasma cell differentiation, we perform differential polyadenylation analysis between plasma cells and B cells. We used the same linear modeling framework as previously described, and used donor identity as a covariate in place of gRNA identity. using linear models with covariates for donor and cell type label. We used a q-value threshold of 1% and required that average polyA sites usage differ by at least 5% between naive B cells and plasma cells. For genes where we observed changes in tandem polyadenylation resulting in 3'UTR shortening, we performed GO Enrichment analysis using the enrichGO function in the R package clusterprofiler[26]. The dotplot function was used for visualization of enrichment results.

In Figure 6J, we consider the set of proximal polyA sites that are more frequently used in plasma cells compared to naive B cells. In each cell, we average the polyA residual across all of these sites, obtaining a single number ('module score') that effectively summarizes the degree of 3' UTR shortening observed within a single cell. We visualize the distribution of these scores across cell types, and then defined two populations of plasma cells based on the median value of this score (Figure 6J). We then performed differential gene expression analysis between the two plasma populations (using logistic regression with donor as a covariate), and performed GO Enrichment analysis on the results using clusterprofiler.
