## Supplementary Figures for "CPA-Perturb-seq: Multiplexed single-cell characterization of alternative polyadenylation regulators"

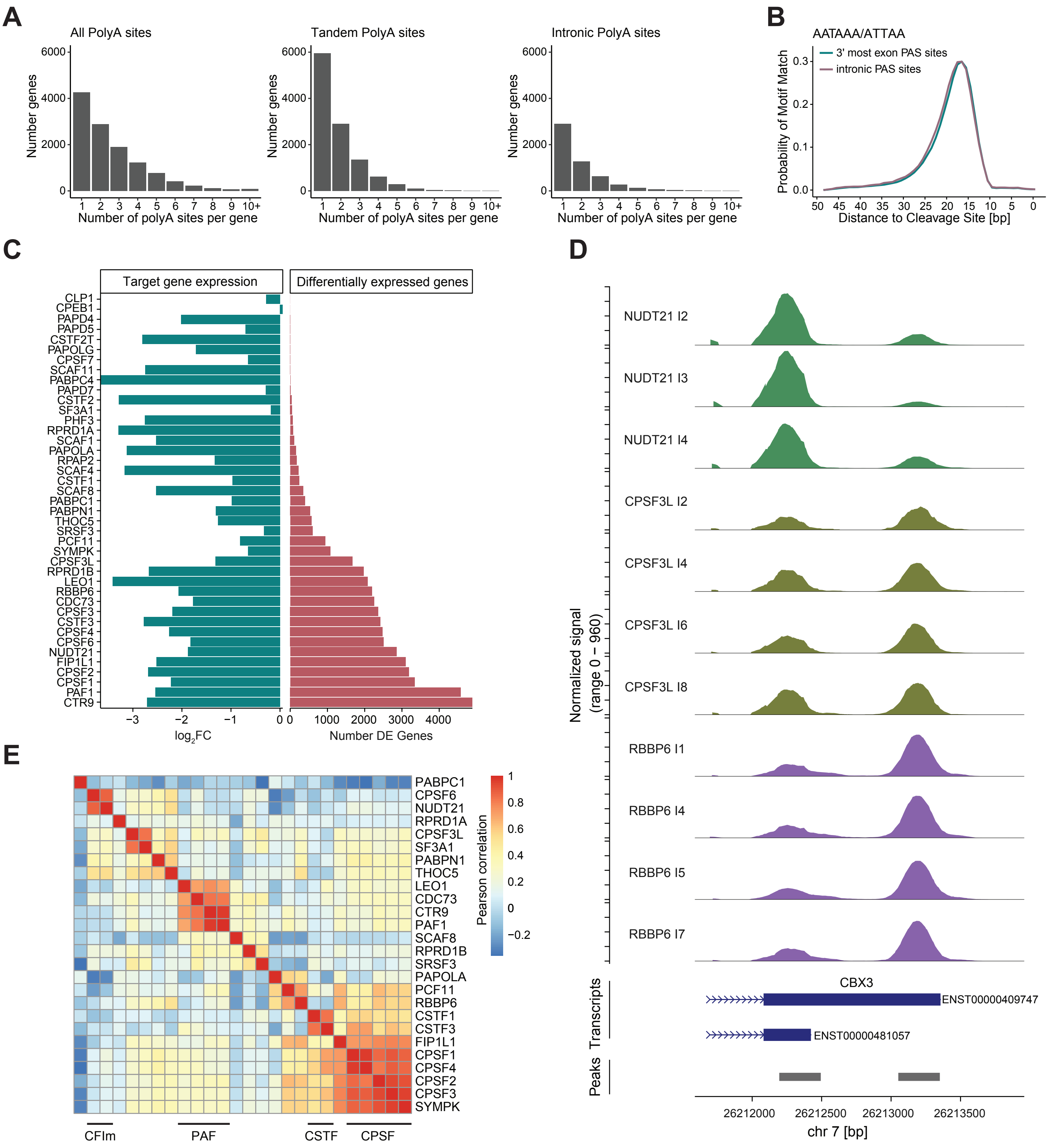


**Supplementary Figure 1: CPA-Perturb-seq dataset**

**(A)** Histogram showing the number of identified polyA sites (after intersection with polyAdbv3) per gene, in the CPA-Perturb-seq dataset. **(B)** Positional preference of the canonical cleavage motives AATAAA and ATTAAA relative to the detected polyA sites in terminal exons and introns. **(C)** For each of the 42 perturbations, the log2FC of the target gene observed in the Perturb-seq data, and the number of differentially expressed genes. In cases where either no or few DE genes were detected, Mixscape will classify all cells as 'non-perturbed'. This can occur in cases where the target perturbation was unsuccessful (i.e. CPEB1), but also in cases where the target gene was downregulated but no global effect was observed (i.e. PABC4). **(D)** Read coverage plot at the CBX3 locus. Each track represents a pseudobulk average of cells, grouped by their perturbation and separated by their individual gRNA. We observe concordant perturbation responses across multiple independent gRNA. **(E)** Hierarchical clustering of the polyA site/regulator count matrix. Each column represents pseudobulk average of single-cells that received the same target gene, after subtracting the pseudobulk average of NT control cells.

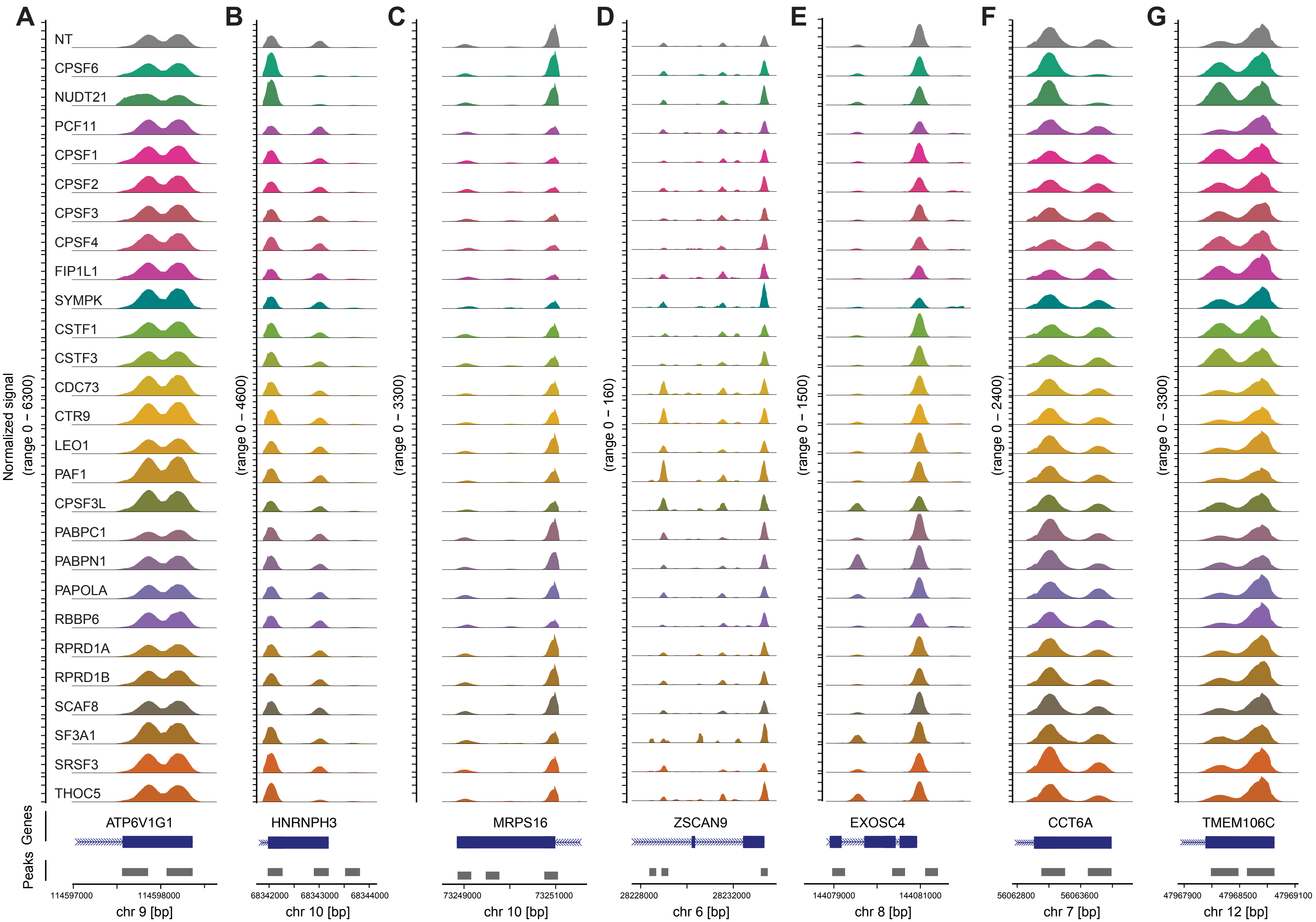


**Supplementary Figure 2: Representative read coverage plots in CPA-Perturb-Seq dataset**

**(A-G)** Read coverage plots depicting the differential use of alternative polyA sites at representative loci. Each track represents a pseudobulk average of cells, grouped by their perturbation. (A to G). Loci were selected based on read coverage plots shown in Figure 1-5, and show data for all 26 regulators and NT control cells. At the ATP6V1G1 locus (A), we observed changes in total gene abundance, rather than relative polyA site usage, across regulators.

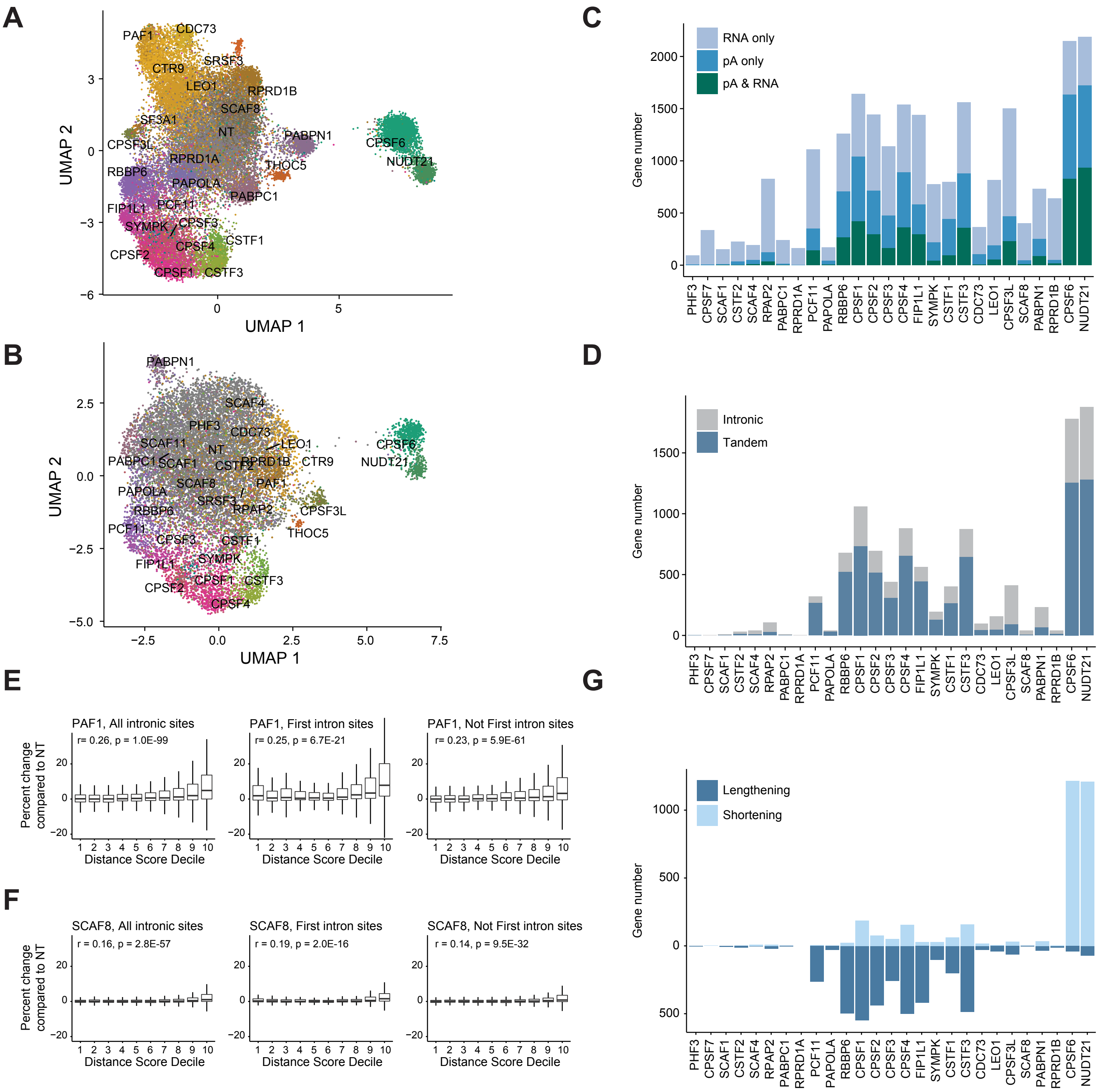


**Supplementary Figure 3: Perturbation-driven changes in tandem and intronic polyadenylation**
**(A-B)** UMAP visualization of (A) HEK293FT cells and (B) K562 cells profiled via CPA-Perturb-Seq. Cells are colored by target gene identity. Visualization was computed based on a linear discriminant analysis (LDA) of polyA-residuals computed on tandem UTR sites. **(C-D)** Same as Figure 3A-B, but barplots show results from the K562 dataset. (E) Percent change (y-axis) in polyA site usage of PAF1 perturbed cells compared to NT control cells for intronic polyA sites as a function of distance to the next polyA site. Intronic polyA sites are binned into deciles (x-axis) for all sites (left), sites within the first intron (middle), and sites not located within the first intron (right). The distance to the next polyA site correlates with the response of an intronic site to PAF1 perturbation. Pearson correlation is computed between the distance (log-scale) and percent change. **(F)** Same as (E), but for SCAF8. **(G)** Same as Figure 3F, but for K562 cells.

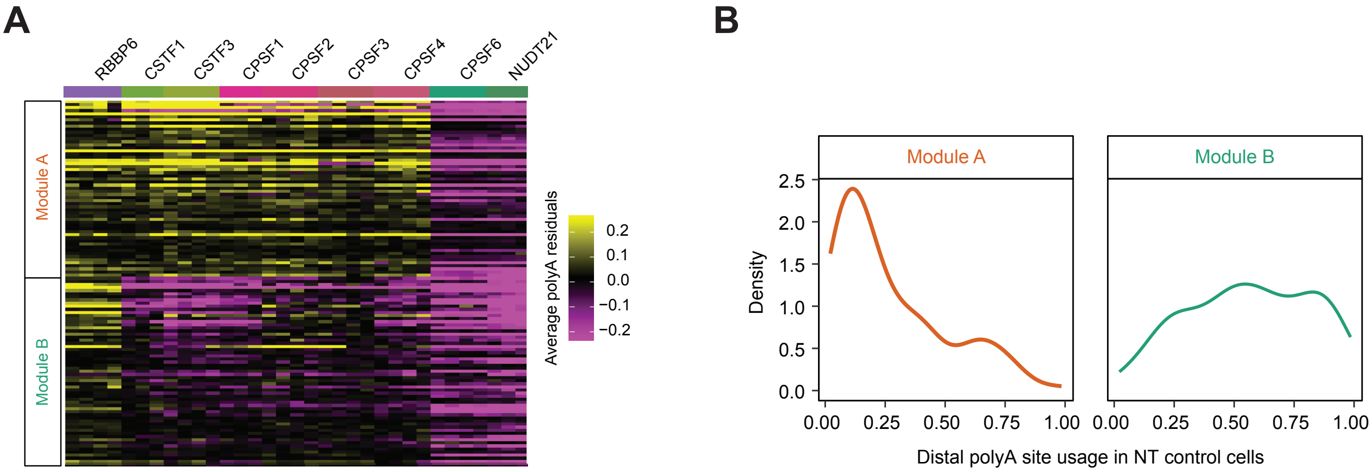


**Supplementary Figure 4: Reproducible APA modules in K562 cells**

**(A)** Same as Figure 4C, heatmap showing K562 polyA residuals for distal sites in Module A genes and Module B genes. Identity and order of polyA sites shown in the heatmap is the same as in Figure 4C. **(B)** Same as Figure 4F, density plot of distal site usage in NT cells for genes belonging to Module A (left) versus Module B (right) in the K562 dataset. Genes in Module A tend to use the proximal site, while genes in Module B tend to use the distal site. Modules were defined in the HEK293 dataset, but the NT distal site usage is plotted for K562 cells.

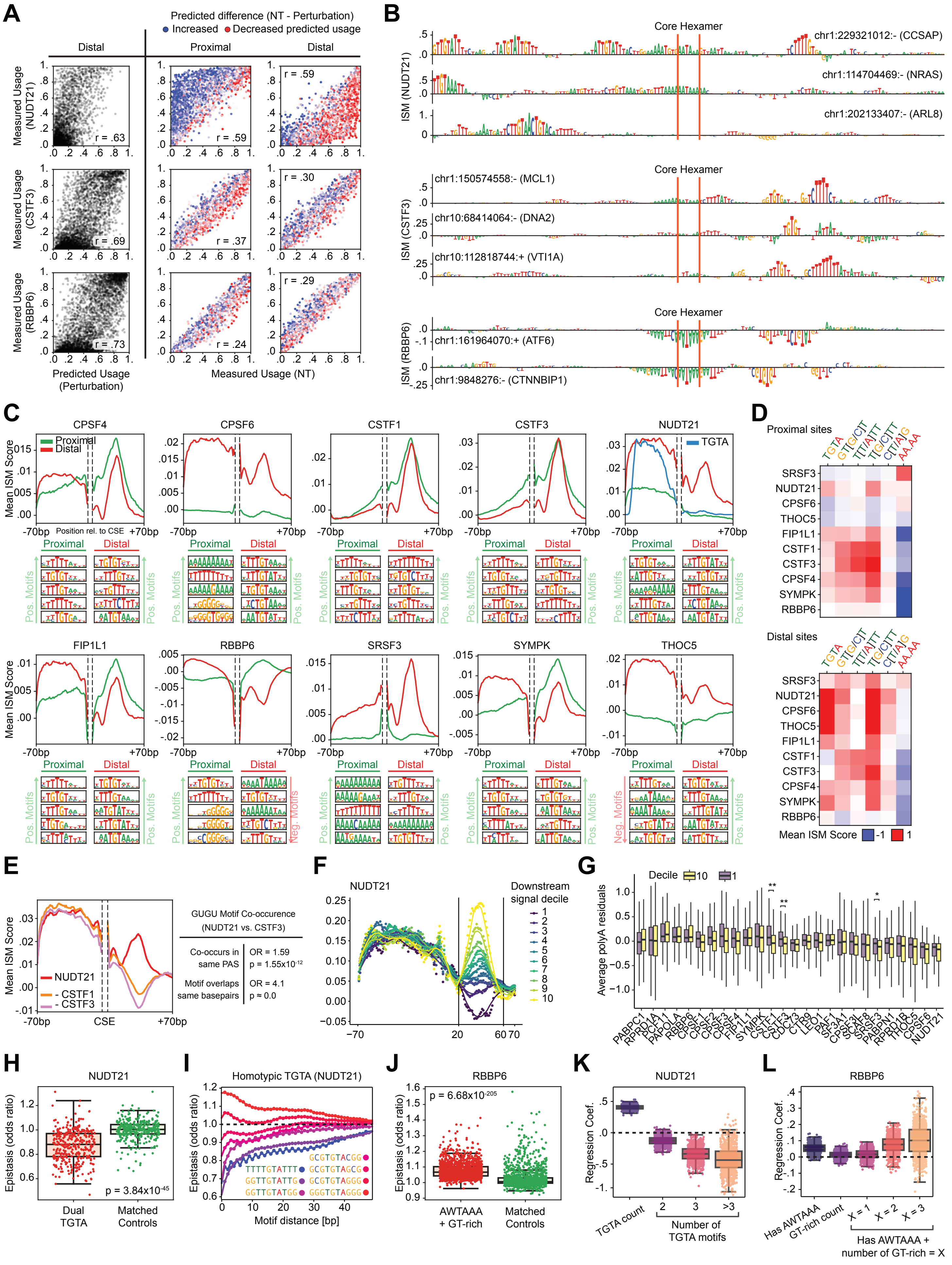

A
Predicted difference (NT - Perturbation)
Increased
Decreased predicted usage
Distal
Proximal
Distal
Measured Usage (NUDT21)
Measured Usage (CSTF3)
Measured Usage (RBBP6)
r = .63
r = .59
r = .59
r = .69
r = .37
r = .30
r = .73
r = .24
r = .29
Predicted Usage (Perturbation)
Measured Usage (NT)
B
Core Hexamer
ISM (NUDT21)
chr1:229321012:- (CCSAP)
chr1:114704469:- (NRAS)
chr1:202133407:- (ARL8)
ISM (CSTF3)
chr1:150574558:- (MCL1)
chr10:68414064:- (DNA2)
chr10:112818744:+ (VTI1A)
ISM (RBBP6)
chr1:161964070:+ (ATF6)
chr1:9848276:- (CTNNBIP1)
C
CPSF4
CPSF6
CSTF1
CSTF3
NUDT21
Proximal
Distal
TGTA
Mean ISM Score
Position rel. to CSE
-70bp
+70bp
Pos. Motifs
FIP1L1
RBBP6
SRSF3
SYMPK
THOC5
Neg. Motifs
D
Proximal sites
Distal sites
TGTA
GT(G/C)T
TT(T/A)TT
T(G/C)TT
C(T/A)G
AA.AA
SRSF3
NUDT21
CPSF6
THOC5
FIP1L1
CSTF1
CSTF3
CPSF4
SYMPK
RBBP6
Mean ISM Score
-1
1
E
Mean ISM Score
NUDT21
- CSTF1
- CSTF3
-70bp
CSE
+70bp
GUGU Motif Co-occurence (NUDT21 vs. CSTF3)
Co-occurs in same PAS
OR = 1.59
p = 1.55x10^-12
Motif overlaps same basepairs
OR = 4.1
p ≈ 0.0
F
NUDT21
Downstream signal decile
1
2
3
4
5
6
7
8
9
10
-70
20
60
70
G
Decile
10
1
Average polyA residuals
**
**
*
PABPC1
RPRD1A
PCF11
PAPOLA
RBBP6
CPSF1
CPSF2
CPSF3
CPSF4
FIP1L1
SYMPK
CSTF1
CSTF3
CDC73
CTR9
LEO1
PAF1
SF3A1
CPSF3L
SCAF8
SRSF3
PABPN1
RPRD1B
THOC5
CPSF6
NUDT21
H
NUDT21
Epistasis (odds ratio)
p = 3.84x10^-45
Dual TGTA
Matched Controls
I
Homotypic TGTA (NUDT21)
Epistasis (odds ratio)
GCGTGTACGG
TTTTGTATTT
GCGTGTAGCG
GGTTGTATTG
GCGTGTAGGG
GGTTGTATGG
GGGTGTAGGG
Motif distance [bp]
J
RBBP6
p = 6.68x10^-205
Epistasis (odds ratio)
AWTAAA + GT-rich
Matched Controls
K
NUDT21
Regression Coef.
TGTA count
2
3
>3
Number of TGTA motifs
L
RBBP6
Regression Coef.
Has AWTAAA
GT-rich count
X = 1
X = 2
X = 3
Has AWTAAA + number of GT-rich = X

**Supplementary Figure 5: Evaluation and interpretation of APARENT-Perturb**

**(A)** (Left) Predicted vs. measured distal polyA site usage in three perturbation conditions (NUDT21, CSTF3 and RBBP6). (Right) Predicted vs. measured difference in proximal or distal polyA site usage with respect to the non-targeting (NT) condition. Only predictions from the held-out sets of the cross-validation procedure are included. **(B)** Same as in Figure 5C, but for additional loci. **(C)** Average residual attribution scores for all 10 learned perturbations, separated by proximal and distal polyA sites. The corresponding top 5 MoDISco motifs are shown below each plot for proximal and distal sites. These motifs are all associated with positive contribution scores (i.e., they increase perturbation magnitude) except for distal motifs in RBBP6 and proximal motifs in THOC5; these motifs are associated with negative contribution scores (they decrease perturbation magnitude). **(D)** Average attribution scores of specific classes of MoDISco motif hits, grouped by perturbation. Red = the motif class is associated with an increase in perturbation magnitude. Blue = the motif class is associated with a decrease in perturbation magnitude. **(E)** Co-occurrence of sequence features with high attribution scores in both the NUDT21 and CSTF1/3 perturbations. (Left) Average attribution scores in the NUDT21 perturbation for distal pA signals (red) and average difference in attribution scores between the NUDT21 perturbation and CSTF1 (orange) or CSTF3 (purple). (Right) Co-occurrence analysis of MoDISco motif hits matching U/G-rich motifs in the NUDT21 and CSTF3 perturbations (odds ratios and p-values calculated with Fisher's exact test). **(F)** Same as Figure 5D, but showing all distal polyA sites that respond to NUDT21 perturbation. PolyA sites are split into deciles based on the model-assigned importance scores for the downstream element region (DSE; denoted by vertical lines). For Decile 1 sites, the DSE is associated with positive attribution scores for NUDT21, while Decile 10 sites have negative contribution scores. **(G)** Average polyA residuals (scaled by perturbation), for regulators in the CPA-Perturb-seq dataset, for polyA sites in Decile 1 vs. Decile 10. These deciles were defined by the NUDT21 perturbation model, but also respond differently to CSTF perturbation. * denotes p-value <0.05 in t-test comparing residuals in Decile 1 vs Decile 10, and ** denotes p-value <0.001. **(H)** Combinatorial ablation of UGUA motifs in distal polyA signals with exactly two wildtype UGUA motifs in the upstream region, which indicates an overall subadditive interaction Epistasis odds ratio (y-axis) estimated by exponentiating the difference in predicted perturbation log odds in the NUDT21 condition when replacing both motifs with random sequence and the sum of log odds when replacing one motif with random sequence at a time. **(I)** Motif insertion simulation for dual UGUA motifs with varying flanking nucleotide context. The y-axis corresponds to the exponentiated difference between the predicted perturbation log-odds in the NUDT21 condition when inserting both motifs and when inserting each motif one at a time. **(J)** Combinatorial ablation of canonical core hexamer motifs and downstream U/G-rich motifs. The U/G-rich motif is replaced with random sequence, while the core hexamer is replaced with a randomly chosen single-nucleotide mutated hexamer (in order to stay in distribution). **(K)** Linear regression coefficients of count features and combinatorial indicator variables when fitting the regression model on distal perturbation log odds ratios in the NUDT21 condition with respect to NT. Distributions of coefficients were generated from 1000-fold bootstrapping. **(L)** Regression coefficient analysis when fitting on RBBP6 perturbation log odds ratios.

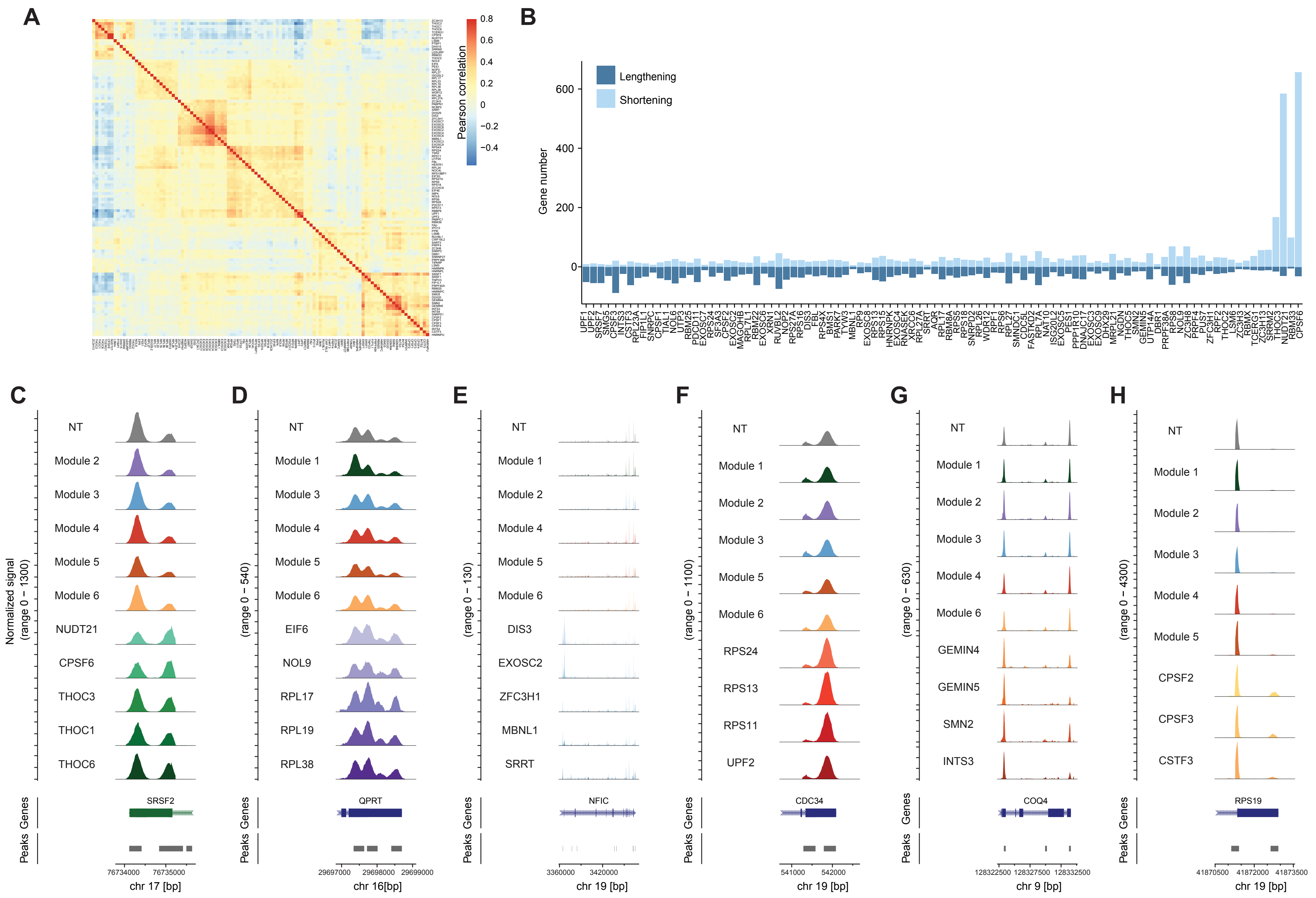


**Supplementary Figure 6: Identifying regulators of alternative polyadenylation from genome-wide Perturb-seq**
**(A)** Same as Figure 6C. Axes show all gene names in the correlation matrix. **(B)** Number of genes with significant changes in tandem polyA site usage in GWPS dataset, classified by 3'UTR shortening or 3'UTR lengthening. **(C-H)** Coverage plots depicting the differential sequencing read coverage and use of alternative polyA sites for representative genes in each module (C to H for Module 1 to 6). Pseudobulk track for NT control cells, and genes present in other modules, are shown for contrast.

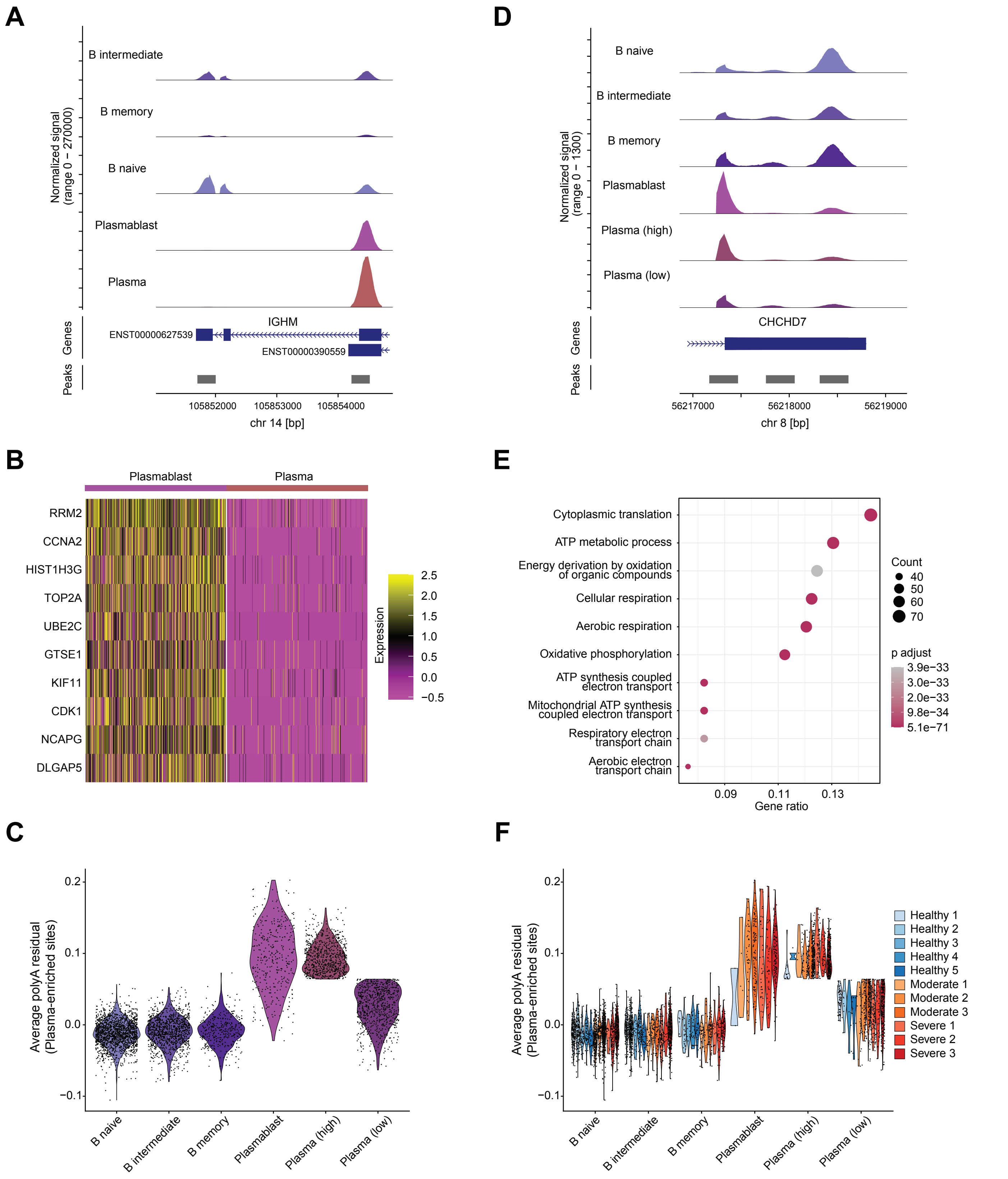


**Supplementary Figure 7: Global shortening of 3'UTR length in Plasma cells and Plasmablasts.**
**(A)** Coverage plot depicting the differential sequencing read coverage and use of alternative polyA sites at the IGHM locus. Each track represents a pseudobulk average of different B and plasma cell subsets. **(B)** Top 10 differentially expressed genes comparing Plasmablasts to Plasma cells. **(C)** Same as Figure 6J, but non-cycling plasma cells were split into two groups (plasma_high, plasma_low), based on the average polyA residual score for proximal plasma-enriched sites. **(D)** Same as Figure 6I, showing the heterogeneity in relative polyA site usage at the CHCHD7 locus across different populations of B and plasma cells. **(E)** Gene ontology analysis of genes more highly expressed in plasma_high compared to plasma_low cells, as indicated in (C). **(F)** Same as in (C), but separated by individual donor. The relationship between cell subset and 3' UTR length is conserved across donors and disease states.
